# A genetically encoded fluorescent sensor for rapid and specific *in vivo* detection of norepinephrine

**DOI:** 10.1101/449546

**Authors:** Jiesi Feng, Changmei Zhang, Julieta Lischinsky, Miao Jing, Jingheng Zhou, Huan Wang, Yajun Zhang, Ao Dong, Zhaofa Wu, Hao Wu, Weiyu Chen, Peng Zhang, Jing Zou, S. Andrew Hires, J. Julius Zhu, Guohong Cui, Dayu Lin, Jiulin Du, Yulong Li

## Abstract

Norepinephrine (NE) and epinephrine (Epi), two key biogenic monoamine neurotransmitters, are involved in a wide range of physiological processes. However, their precise dynamics and regulation remain poorly characterized, in part due to limitations of available techniques for measuring these molecules *in vivo*. Here, we developed a family of <u>G</u>PC<u>R A</u>ctivation-<u>B</u>ased <u>NE</u>/Epi (GRAB_NE_) sensors with a 230% peak ΔF/F_0_ response to NE, good photostability, nanomolar-to-micromolar sensitivities, sub-second rapid kinetics, high specificity to NE vs. dopamine. Viral- or transgenic- mediated expression of GRAB_NE_ sensors were able to detect electrical-stimulation evoked NE release in the locus coeruleus (LC) of mouse brain slices, looming-evoked NE release in the midbrain of live zebrafish, as well as optogenetically and behaviorally triggered NE release in the LC and hypothalamus of freely moving mice. Thus, GRAB_NE_ sensors are a robust tool for rapid and specific monitoring of *in vivo* NE/Epi transmission in both physiological and pathological processes.

## Introduction

Both norepinephrine (NE) and epinephrine (Epi) are key monoamine neurotransmitters in the central nervous systems and peripheral organs of vertebrate organisms. These transmitters play an important role in a plethora of physiological processes, allowing the organism to cope with its ever-changing internal and external environment. In the brain, NE is synthesized primarily in the locus coeruleus (LC), a small yet powerful nucleus located in the pons. Noradrenergic LC neurons project throughout the brain and exert a wide range of effects, including processing sensory information (Berridge and Waterhouse, 2003), regulating the sleep-wake/arousal state (Berridge et al., 2012), and mediating attentional function (Bast et al., 2018). Blocking noradrenergic transmission causes impaired cognition and arousal and is closely correlated with a variety of psychiatric conditions and neurodegenerative diseases, including stress (Chrousos, 2009), anxiety (Goddard et al., 2010), depression (Moret and Briley, 2011), attention-deficit hyperactivity disorder (ADHD) (Berridge and Spencer, 2016), and Parkinson’s disease (PD) (Espay et al., 2014). In the sympathetic nervous system, both NE and Epi play a role in regulating heart function (Brodde et al., 2001) and blood pressure (Zimmerman, 1981).

Despite their clear importance in a wide range of physiological processes, the spatial and temporal dynamics of NE and Epi in complex organs (*e.g.* the vertebrate brain) are poorly understood at the *in vivo* level due to limitations associated with current detection methods. Classic detection methods such as microdialysis-coupled biochemical analysis (Bito et al., 1966; Justice, 1993; Watson et al., 2006) have low temporal resolution, requiring a relatively long time (typically 5 min/collection) and complex sampling procedures, thereby limiting the ability to accurately measure the dynamics of noradrenergic activity in the physiological state (Chefer et al., 2009). Recent improvements in microdialysis—in particular, the introduction of the nano-LC-microdialysis method (Lee et al., 2008; Olive et al., 2000)—have significantly increased detection sensitivity; however, this approach is still limited by a relatively slow sampling rate (on the order of several minutes). On the other hand, electrochemical detection techniques based on measuring currents generated by the oxidation of NE/Epi (Bruns, 2004; Park et al., 2009; Robinson et al., 2008; Zhou and Misler, 1995) provide nanomolar sensitivity and millisecond temporal resolution; however, their inability to distinguish NE and Epi from other monoamine neurotransmitters— particularly dopamine (Robinson et al., 2003)—presents a significant physiological limitation with respect to measuring noradrenergic/adrenergic transmission both in *ex vivo* tissue preparations and *in vivo*. In addition, both microdialysis-based and electrochemical techniques are designed to detect volume-averaged NE/Epi levels in the extracellular fluid and therefore cannot provide cell type–specific or subcellular information.

Real-time imaging of NE dynamics would provide an ideal means to non-invasively track NE with high spatiotemporal resolution. A recent innovation in real-time imaging, the cell-based reporters known as CNiFERs (Muller et al., 2014), converts an extracellular NE signal into an intracellular calcium signal that can be measured using highly sensitive fluorescence imaging. However, CNiFERs require implantation of exogenous cells and can report only volume transmission of NE/Epi. By contrast, genetically encoded sensors, in theory, circumvent the above-mentioned limitations to provide fast, clear, non-invasive, cell type–specific reporting of NE/Epi dynamics. In practice, all genetically encoded NE sensors developed to date have poor signal-to-noise ratio and narrow dynamic range (*e.g.*, a <10% change in FRET ratio under optimal conditions) (Nakanishi et al., 2006; Vilardaga et al., 2003; Wang et al., 2018b), thus limiting their applicability, particularly in *in vivo* applications.

To overcome these limitations, we developed a series of genetically encoded single-wavelength fluorescent GRAB_NE_ sensors with rapid kinetics, a ΔF/F_0_ dynamic range of ∽200%, and EGFP-comparable spectra, brightness, and photostability. Here, we showcase the wide applicability of our GRAB_NE_ sensors using a number of *in vitro* and *in vivo* preparations. In every application tested, the GRAB_NE_ sensors readily reported robust, chemical-specific NE signals. Thus, our GRAB_NE_ sensors provide a powerful imaging-based probe for measuring the cell-specific regulation of adrenergic/noradrenergic transmission under a wide range of physiological and pathological conditions.

## Results

### Development and characterization of GRAB_NE_ sensors

Inspired by the structure (Rasmussen et al., 2011a; Rasmussen et al., 2011b) and working mechanism (Chung et al., 2011; Manglik et al., 2015; Nygaard et al., 2013) of the β2 adrenergic G protein–coupled receptor (GPCR), we exploited the conformational change between the fifth and sixth transmembrane domains (TM5 and TM6, respectively) upon ligand binding to modulate the brightness of an attached fluorescent protein. Building upon the successful strategy of generating GPCR activation-based sensors for acetylcholine (GACh) (Jing et al., 2018) and dopamine (GRAB_DA_) (Sun et al., 2018), we first systematically screened human adrenergic receptors as a possible scaffold. We inserted circular permutated EGFP (cpEGFP) into the third intracellular loop domain (ICL3) of three α-adrenergic receptors (α1DR, α2AR, and α2BR) and two β-adrenergic receptors (β2R and β3R) (Fig. 1A). Among these five constructs, we found that α2AR-cpEGFP had the best membrane trafficking, indicated by its high colocalization with membrane-targeted RFP (Fig. S1); we therefore selected this construct as the scaffold for further screening.

**Figure 1.**
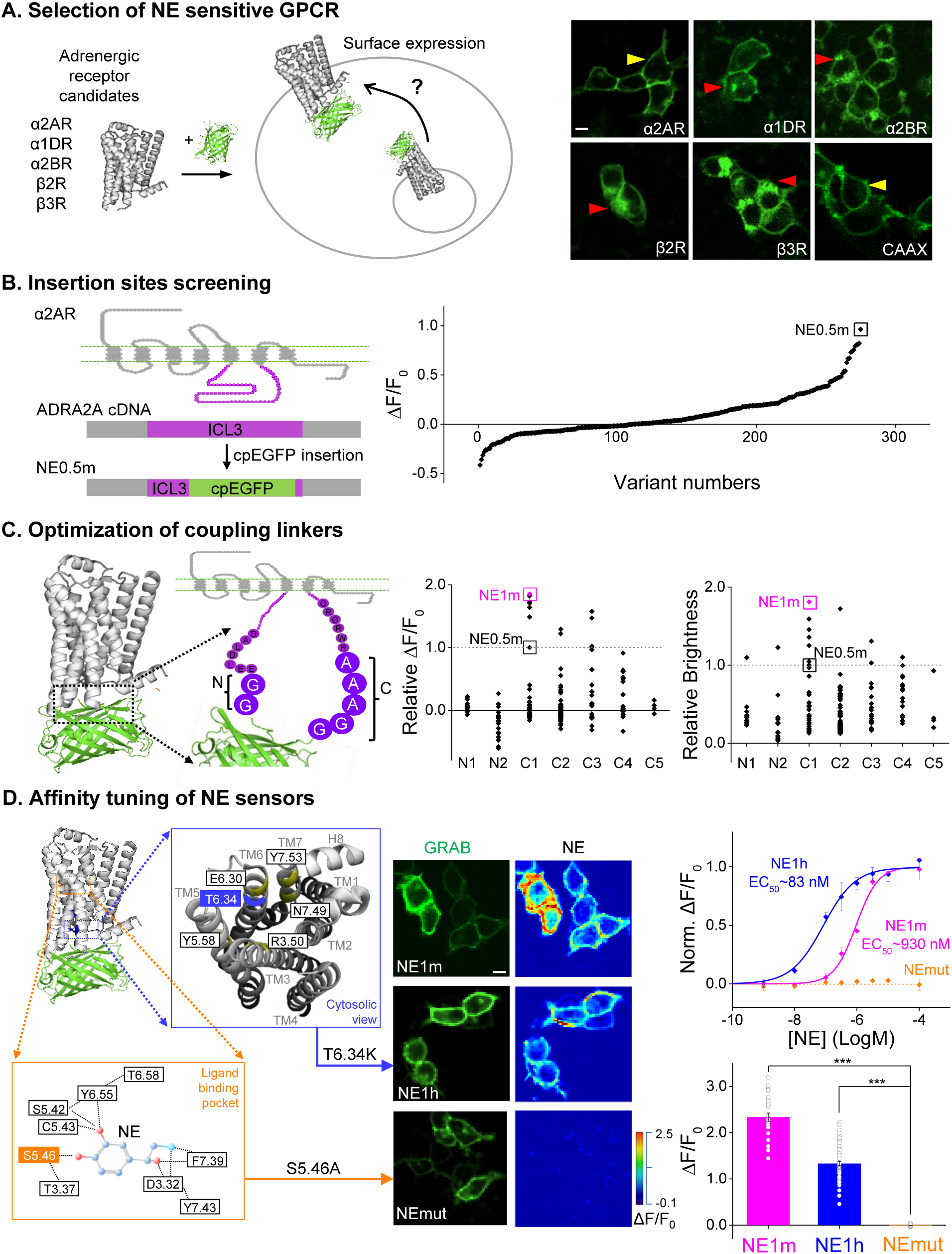
Figure 1. Design and optimization of genetically encoded NE sensors. (**A**) Selection of a candidate sensor scaffold by screening several NE-binding GPCRs. Shown at the right are example images of the indicated chimeric GPCR-cpEGFP candidates expressed in HEK293T cells. Yellow arrows indicate robust membrane trafficking, and red arrows indicate impaired membrane trafficking. See also Figure S1. (B) Identification of the most responsive NE sensor, NE0.5m (indicated by the black square) by screening the cpEGFP insertion site in ICL3 of the α2AR. ΔF/F_0_ refers to the peak change in fluorescence intensity in response to 100 μM NE. (C) Optimizing the GRAB_NE_ sensors by mutational screening of the insertion linker. NE0.5m was used as a template, and the indicated amino acids on N-terminal and C-terminal sides of the cpEGFP insert were mutated individually. Sensor NE1m (indicated by the pink squares) was identified due to having the strongest response (ΔF/F_0_) and brightness relative to the original NE0.5m sensor (indicated by the dashed line at 1.0). (D) Tuning the sensor’s affinity for NE by introducing mutations in the GPCR. Magnified views of the ligand-binding pocket view from the cytosol are shown; key residues involved in ligand binding and inducing a conformational change upon ligand binding are indicated. The middle panel shows example images of HEK293T cells expressing the indicated GRAB_NE_ sensors; EGFP fluorescence is shown in the left column, and the fluorescence response in the presence of 100 μM NE is shown in the right column. Shown at the right are the normalized dose-response curves for the three GRAB_NE_ sensors, with C_50_ values (top), and the average fluorescence change in response to 100 μM NE (bottom); n = 21-67 cells from 3-5 cultures for each sensor. The scale bars in (A) and (D) represent 10 μm. \*\*\**p* < 0.001 (Student’s *t*-test).

The length of the linker surrounding the cpEGFP moiety inserted in G-GECO (Zhao et al., 2011), GCaMP (Akerboom et al., 2012), GACh (Jing et al., 2018), and GRAB_DA_ (Sun et al., 2018) can affect the fluorescence response of cpEGFP-based indicators. Thus, as the next step, we systematically truncated the linker which starts with the entire flexible ICL3 of α2AR surrounding cpEGFP (Fig. 1B). We initially screened 275 linker-length variant proteins and identified a sensor (GRAB_NE0.5m_) with a modest response to NE (Fig. 1B, right). From this scaffold, we performed a random mutation screening of seven amino acids (AAs) in close proximity to the cpEGFP moiety; two of these AAs are on the N-terminal side of cpEGFP, and the remaining five are on the C-terminal side of cpEGFP (Fig. 1C). From approximately 200 mutant versions of GRAB_NE0.5m_, we found that GRAB_NE1m_—which contains a glycine-to-threonine mutation at position C1—provided the best performance with respect to ΔF/F_0_ and brightness (Fig. 1C, middle and right).

Next, we expressed GRAB_NE1m_ in HEK293T cells and applied NE in a range of concentrations. NE induced a fluorescence change in GRAB_NE1m_-expressing cells in a dose-dependent manner, with an EC_50_ of 0.93 μM and a maximum ΔF/F_0_ of approximately 230% in response to a saturating concentration of NE (100 μM) (Fig. 1D, middle and right). We also introduced mutations in α2AR in order to increase its sensitivity at detecting NE. We found that a single T6.34K point mutation (Ren et al., 1993)—which is close to the highly conserved E6.30 site—resulted in a 10-fold increase in sensitivity (EC_50_ ∽83 nM) to NE compared with GRAB_NE1m_; this sensor, which we call GRAB_NE1h_, has a maximum ΔF/F_0_ of ∽130% in response to 100 μM NE. As a control, we also generated GRAB_NEmut_, which has the mutation S5.46A at the putative ligand-binding pocket and therefore is unable to bind NE (Fig. 1D); this control sensor has similar brightness and membrane trafficking (Fig. S1 and S2A), but does not respond to NE even at 100 μM (Fig. 1D, middle and right).

To examine whether our GRAB_NE_ sensors can capture the rapid dynamic properties of NE signaling, including its release, recycling, and degradation, we bathed GRAB_NE1h_-expressing HEK293T cells in a solution containing NPEC-caged NE; a focused spot of 405-nm light was applied to locally uncage NE by photolysis (Fig. 2A). Transient photolysis induced a robust increase in fluorescence in GRAB_NE1h_-expressing cells (mean on time constant 137ms, single exponential fit), which was blocked by application of the α2-adrenergic receptor antagonist yohimbine (Fig. 2B,C). To characterize both the on and off rates (τ_on_ and τ_off_, respectively) of the GRAB_NE_ sensors, we locally applied various compounds to GRAB_NE_-expressing cells using rapid perfusion and measured the fluorescence response using high-speed line scanning (Fig. 2D,E). The average delay intrinsic to the perfusion system (measured by fitting the fluorescence increase in the co-applied red fluorescent dye Alexa 568) was 34 ms (Fig. 2F). Fitting the fluorescence change in each sensor with a single exponential function yielded an average τ_on_ of 72 and 36 ms for GRAB_NE1m_ and GRAB_NE1h_, respectively, and an average τ_off_ of 680 and 1890 ms for GRAB_NE1m_ and GRAB_NE1h_, respectively (Fig. 2E,F). The faster on-rate and slower off-rate of GRAB_NE1h_ compared to GRAB_NE1m_ is consistent with its relatively higher affinity for NE.

**Figure 2.**
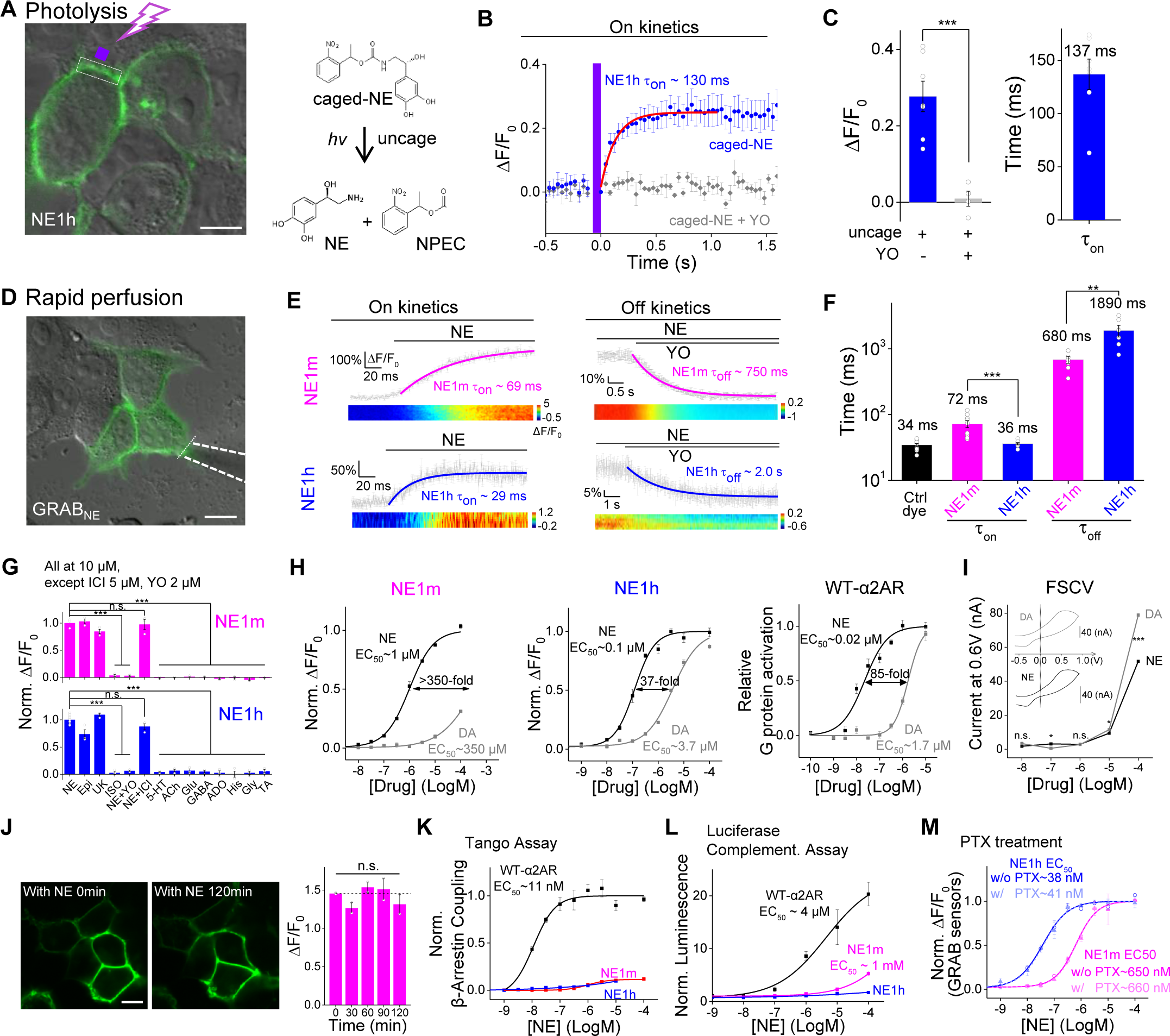
Figure 2. Characterization of GRAB_NE_ sensors in cultured cells. (**A-C**) HEK293T cells were loaded with NPEC-NE, which was uncaged by photolysis with a pulse of 405-nm light. Uncaging caused a rapid increase in GRAB_NE1h_ fluorescence, which was blocked in the presence of 10 μM yohimbine (YO). The data in A represent 3 trials each, and the data in C represent 7 cells from 3 cultures. The white dotted square indicates the image region and the purple square indicates the illumination region. (**D-F**) NE was applied to HEK293T cells expressing GRAB_NE1m_ or GRAB_NE1h_ to measure τ_on_. Yohimbine (YO) was then applied in order to measure τ_off_; The white dotted line indicates the line-scanning region. n ≥ 6 cells from 6 cultures. (**G**) The indicated compounds were applied to GRAB_NE1m_ and GRAB_NE1h_, and the change in fluorescence relative to NE is plotted. (**H**) Dose-response curves for GRAB_NE1m_, GRAB_NE1h_, and wild-type α2AR for NE and DA, with EC_50_ values shown; n ≥ 3 wells with 100-300 cells each. (**I**) Fast-scan cyclic voltammetry measurements in response to increasing concentrations of NE and DA. The insets show exemplar cyclic voltammograms of NE and DA at 100 μM, with peak current occurring at ∽0.6 V. (**J**) Time course of ΔF/F_0_ for GRAB_NE_ sensors measured over a 2-h time frame; note that the fluorescent signal remained at the cell surface even after 180 min, indicating no measurable internalization or desensitization. n = 3 wells with 100-300 cells each. (**K**) A TANGO assay was performed in order to measure β-arrestin–mediated signaling by GRAB_NE1m_, GRAB_NE1h_, and wild-type α2AR in the presence of increasing concentrations of NE; n = 4 wells with ≥10^5^ cells each. (**L,M**) GRAB_NE_ sensors do not couple to downstream G protein signaling pathways. Wild-type α2AR, but not GRAB_NE1m_ or GRAB_NE1h_, drives Gαi signaling measured using a luciferase complementation assay (L). Disrupting of G protein activation with pertussis toxin does not affect the NE-induced fluorescence change in GRAB_NE1m_ or GRAB_NE1h_ (M). n = 3 wells with ≥10^5^ cells each. The scale bars in (A), (D), and (J) represent 10 μm. \**p* < 0.05, \*\**p* < 0.01, and \*\*\**p* < 0.001; n.s., not significant (Student’s *t*-test).

High ligand specificity is an essential requirement for tools designed to detect structurally similar monoamine-based molecules. Importantly, our GRAB_NE_ sensors, which are based on α2AR, respond to both NE and Epi, but do not respond to other neurotransmitters (Fig. 2G). The sensors also respond to the α2AR agonist brimonidine but not the β2-adrenergic receptor agonist isoprenaline, which indicates receptor-subtype specificity. Moreover, the NE-induced fluorescence increase in GRAB_NE_-expressing cells was blocked by the α-adrenergic receptor antagonist yohimbine, but not the β-adrenergic receptor antagonist ICI 118,551. Additionally, because NE and DA are structurally similar yet functionally distinct, we characterized how our GRAB_NE_ sensors respond to various concentrations of DA and NE. Wild-type α2AR has an 85-fold higher affinity for NE versus DA (Fig. 2H, right); in contrast, GRAB_NE1m_ has a 350-fold higher affinity for NE, whereas GRAB_NE1h_ was similar to the wild-type receptor, with a 37-fold higher affinity for NE (Fig. 2H). In contrast, fast-scan cyclic voltammetry (FSCV) was unable to differentiate between NE and DA, producing a nearly identical response to similar concentrations of NE and DA (Fig. 2I) (Robinson et al., 2003). To test the photostability of our NE sensors, we continuously illuminated GRAB_NE_-expressing HEK293T cells using either 1-photon (confocal) or 2-photon laser microscopy and found that the GRAB_NE_ sensors are more photostable than EGFP under both conditions (Fig. S2C). Taken together, these data suggest that the GRAB_NE_ sensors can be used to measure the dynamic properties of noradrenergic activity with high specificity for NE over other neurotransmitters.

Next, we examined whether our GRAB_NE_ sensors can trigger GPCR-mediated downstream signaling pathways. First, we bathed GRAB_NE1m_-expressing cells in a saturating concentration of NE for 2 h, but found no significant internalization of GRAB_NE1m_ (Fig. 2J). Similarly, we found that both GRAB_NE1m_ and GRAB_NE1h_ lack β-arrestin–mediated signaling, even at the highest concentration of NE tested (Fig. 2K), suggesting that the GRAB_NE_ sensors are not coupled to β-arrestin signaling. In addition, GRAB_NE1m_ and GRAB_NE1h_ had drastically reduced downstream Gi coupling compared to wild-type α2AR, which was measured using a Gi-coupling–dependent luciferase complementation assay (Fig. 2L) (Wan et al., 2018). We also found that G protein activation by GRAB_NE1m_ measured using the highly sensitive TGFα shedding was reduced by about 100-fold compared to the wild-type receptor (Fig. S2B) (Inoue et al., 2012). Finally, blocking G protein activation by treating cells with pertussis toxin (Fig. 2M) had no effect on the fluorescence response of either GRAB_NE1m_ or GRAB_NE1h_, indicating that the fluorescence response of GRAB_NE_ sensors does not require G protein coupling (Rasmussen et al., 2011a). Taken together, these data indicate that GRAB_NE_ sensors can be used to report NE concentration without inadvertently engaging GPCR downstream signaling.

### Characterization of GRAB_NE_ sensors in cultured neurons

The expression, trafficking, and response of proteins can differ considerably between neurons and cell lines (Marvin et al., 2013; Zou et al., 2014). Therefore, to characterize the performance of GRAB_NE_ sensors in neurons, we co-expressed GRAB_NE_ together with several neuronal markers in cultured cortical neurons. Both GRAB_NE1m_ and GRAB_NEmut_ trafficked to the cell membrane and co-localized with the membrane-targeted marker RFP-CAAX (Fig. 3A,B). Upon bath-application of a saturating concentration of NE, GRAB_NE1m_ and GRAB_NE1h_ had a peak ΔF/F_0_ of approximately 230% and 150%, respectively, whereas GRAB_NEmut_ had no response (Fig. 3D,E); these results are similar to our results obtained with HEK293T cells. Moreover, the NE-induced response in GRAB_NE1m_-expressing cells was similar among various subcellular compartments identified by co-expressing GRAB_NE1m_ with either the axonal marker synaptophysin (SYP) or the dendritic marker PSD95 suggesting that GRAB_NE_ sensors enable the detection of NE throughout the neurons (Fig. 3C). Both GRAB_NE1m_- and GRAB_NE1h_-expressing neurons had a dose-dependent fluorescence increase in response to NE, with mean EC_50_ values of 1.9 μM and 93 nM, respectively (Fig. 3F). Consistent with high selectivity for NE, GRAB_NE1m_ and GRAB_NE1h_ have a 1000-fold and 7-fold higher affinity, respectively, for NE versus DA (Fig. 3F). Moreover, GRAB_NE1m_ responded specifically to NE and Epi, but did not respond to several other neurotransmitters and ligands, including the β2-adrenergic receptor agonist isoprenaline, histamine, dopamine, and serotonin (Fig. 3G). Finally, culturing GRAB_NE1m_-expressing neurons in 100 μM NE for one hour did not cause internalization of the sensor, and the fluorescence increase was both stable for the entire hour and blocked completely by the α2-adrenergic receptor antagonist yohimbine (Fig. 3H,I). Thus, our GRAB_NE_ sensors have the necessary affinity and specificity to faithfully measure noradrenergic signaling in neurons.

**Figure 3.**
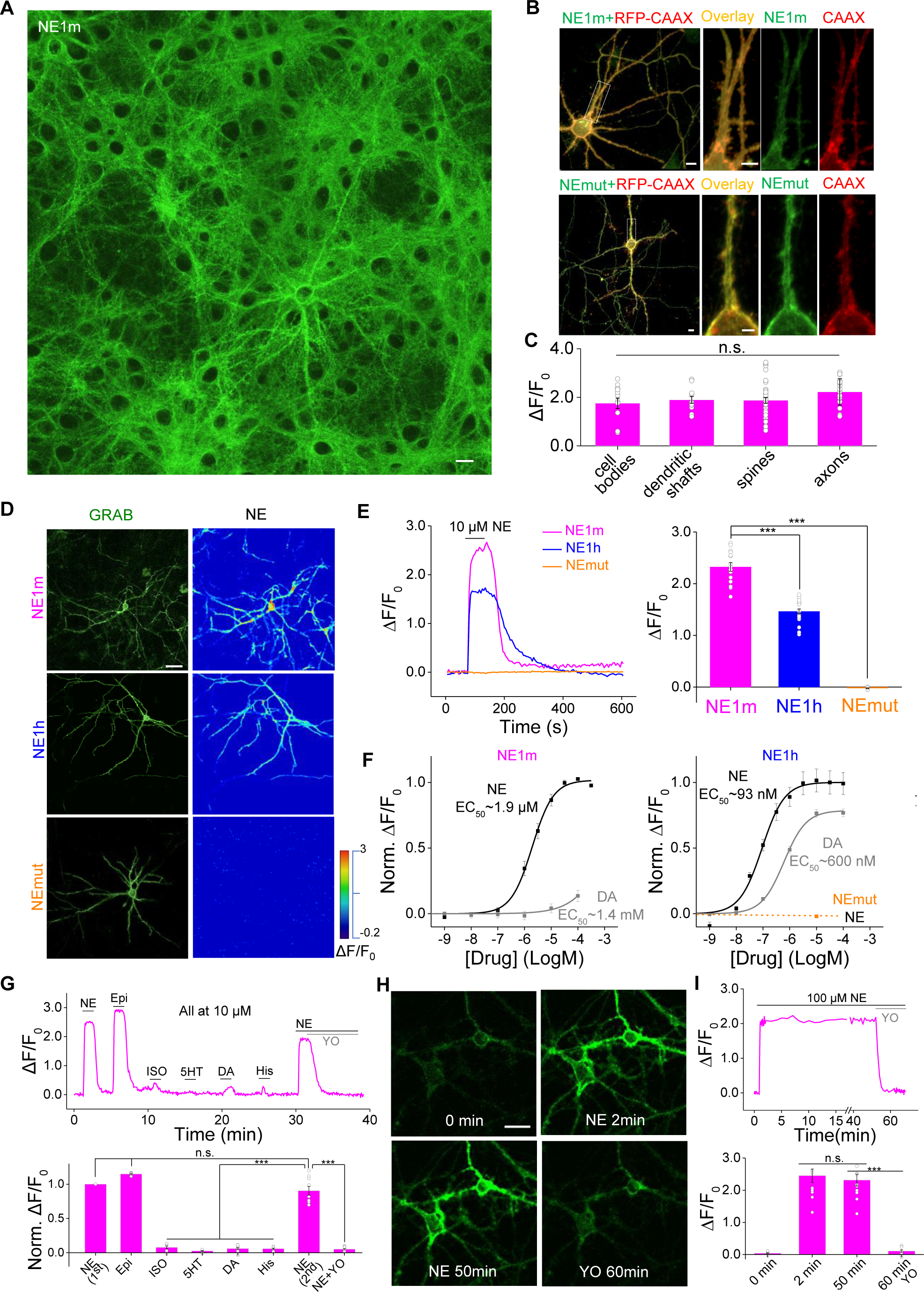
Characterization of GRAB_NE_ sensors in cultured neurons. (**A-C**) GRAB_NE1m_ is expressed in various plasma membrane compartment of cultured neurons. Cultured cortical neurons were co-transfected with GRAB_NE1m_ and RFP-CAAX to label the plasma membrane, and the fluorescence response induced by bath application of NE was measured in the cell body, dendritic shaft and spine, and axon (**C**). n > 10 neurons from 4 cultures. (**D,E**) Cultured cortical neurons expressing GRAB_NE1m_ and GRAB_NE1h_, but not GRAB_NEmut_, respond to application of NE (10 μM). EGFP fluorescence and pseudocolor images depicting the response to NE are shown in (**D**), and the time course and summary of peak ΔF/F_0_ are shown in (**E**). n > 15 neurons from 3 cultures. (**F**) Dose-response curve for GRAB_NE_ sensors expressed in cultured cortical neurons in response to NE and DA. n > 10 neurons from 3 cultures. (**G**) Example trace (top) and summary (bottom) of cultured neurons transfected with GRAB_NE1m_ and treated with the indicated compounds at 10 μM each. n = 9 neurons from 3 cultures. (**H,I**) The fluorescence change in GRAB_NE1m_ induced by 100 μM NE is stable for up to 1 h. Representative images taken at the indicated times are shown in (**H**). An example trace and summary data are shown in (**I**). Where indicated, 10 μM yohimbine (YO) was added. n = 11 neurons from 3 cultures. The scale bars in (**A**) and (**B**) represent 10 μm; the scale bars in (**D**) and (**H**) represent 25 μm. \*\*\**p* < 0.001; n.s., not significant (Student’s *t*-test).

### Characterization of GRAB_NE_ sensors in both cultured and acute brain slices

To further test the GRAB_NE_ sensors *in vitro,* we expressed GRAB_NE1m_ and GRAB_NE1h_ in cultured hippocampal slices using a Sindbis virus expression system (Fig. S3A). In both GRAB_N_E1m-expressing CA1 neurons and GRAB_NE1h_-expressing CA1 neurons, exogenous application of NE in ACSF—but not ACSF alone—evoked a robust increase in fluorescence (Fig. S3B-D). In contrast, NE had no detectable effect on GRAB_N_Emut-expressing neurons (Fig. S3C,D). Application of several α-adrenergic receptor agonists, including epinephrine and brimonidine, also generated a fluorescence increase in GRAB_NE1m_-expressing neurons (Fig. S3C,F), consistent with data obtained using cultured cells. The rise and decay kinetics of the change in fluorescence were second-order, which reflects the integration of the time required to puff the drugs onto the cells and the sensor’s response kinetics (Fig. S3E,G). We also prepared acute hippocampal slices in which GRAB_NE1h_ was expressed using an adeno-associated virus (AAV); in this acute slice preparation, the GRAB_N_E1h-expressing hippocampal neurons are innervated by noradrenergic fibers, which was confirmed by post-hoc staining using an antibody against dopamine beta hydroxylase (Fig. S3H,I). Application of electrical stimuli at 20 Hz for 1 s elicited a robust increase in GRAB_N_E1h fluorescence, and this increase was blocked by the application of yohimbine (Fig. S3J). Consistent with our results obtained using cultured slices, exogenous application of various α-adrenergic receptor agonists, including NE, Epi, and brimonidine— but not the β-adrenergic receptor agonist isoprenaline—evoked a fluorescence increase in GRAB_N_E1h-expressing neurons, and this response was blocked by yohimbine, but not by the β-adrenergic receptor antagonist ICI 118,551 (Fig. S3K).

Next, we examined whether our GRAB_NE_ sensors can be used to monitor the dynamics of endogenous NE. We expressed GRAB_NE1m_ in the locus coeruleus (LC), which contains the majority of adrenergic neurons within the brain (Fig. 4A). Two weeks after AAV injection, we prepared acute brain slices and observed GRAB_NE1m_ expression in the membrane of LC neurons using two-photon microscopy (Fig. 4A). We then used electrical stimuli to evoke the release of endogenous NE in the LC in the acute slices. Applying one or two stimuli did not produce a detectable fluorescence increase in GRAB_NE1m_-expressing neurons; in contrast, applying 10 or more stimuli at 20 Hz caused a progressively stronger response (Fig. 4B). Application of the voltage-activated potassium channel blocker 4-aminopyridine, which increases Ca^2+^ influx during the action potential, significantly increased the fluorescence response, whereas application of Cd^2+^ to block calcium channels abolished the stimulation-induced fluorescence increase (Fig. 4C), consistent with presynaptic NE release being mediated by Ca^2+^ influx. We also performed line-scanning experiments in order to track the kinetics of NE release (Fig. 4D, left). A brief electrical stimulation induced a rapid fluorescence response with a mean τ_on_ and τ_off_ of 37 ms and 600 ms, respectively (Fig. 4D, middle and right). Taken together, these data indicate that GRAB_NE1m_ can be used to monitor the release of endogenous NE in real time.

**Figure 4.**
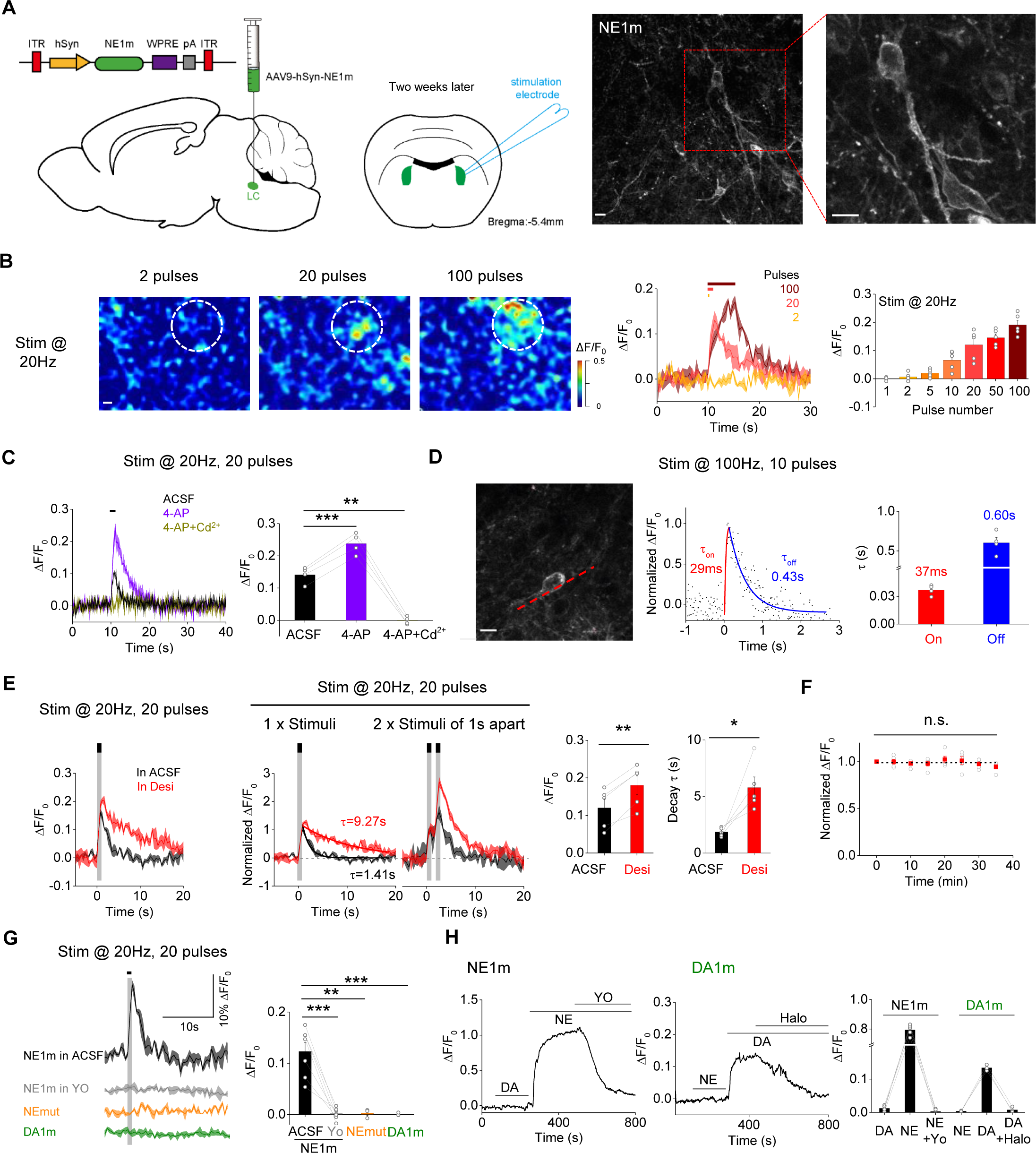
Release of endogenous NE measured in mouse brain slices. **(A) Left**, schematic illustration of the slice experiments. An AAV expressing hSyn-NE1m was injected into the LC; two weeks later, acute brain slices were prepared and used for electric stimulation experiments. **Right**, exemplar 2-photon microscopy images showing the distribution of GRAB_NE1m_ in the plasma membrane of LC neurons. **(B) Left and middle**, representative pseudocolor images and corresponding fluorescence changes in GRAB_NE1m_-expressing neurons in response to 2, 20, and 100 pulses delivered at 20 Hz. The ROI (50-μm diameter) for data analysis is indicated in the images. **Right**, summary of the peak fluorescence change in slices stimulated as indicated; n = 5 slices from 5 mice. **(C)** Exemplar traces and summary data of GRAB_NE1m_-expressing neurons in response to 20 electrical stimuli delivered at 20 Hz in ACSF, 4-AP (100 μM), or 4-AP with Cd^2+^ (100μM); n = 4 slices from 4 mice. (**D**) Kinetic properties of the electrically evoked fluorescence responses in GRAB_NE1m_-expressing LC neurons. **Left**, image showing a GRAB_NE1m_-expressing LC neuron for line scan analysis (red dashed line). **Middle and right**, example trace and summary of the responses elicited in GRAB_NE1m_-expressing neurons before, and after 10 pulses delivered at 100Hz; n = 4 slices from 4 mice. (**E**) The norepinephrine transporter blocker desipramine (Desi, 10 μM; red) increases the effect of electrical stimuli (20 pulses at 20 Hz) or two trains of stimuli with a 1-s interval compared to ACSF (black traces). n = 5 slices from 5 mice. (**F**) The fluorescence response in GRAB_NE1m_-expressing neurons is stable. Eight stimuli (20 pulses at 20 Hz) were applied at 5-min intervals, and the response (normalized to the first train) is plotted against time. n = 5 slices from 5 mice. (**G**) Traces and summary data of the fluorescence response measured in neurons expressing GRAB_NE1m_, GRAB_NEmut_, or GRAB_DA1m_ in response to 20 pulses delivered at 20 Hz in the presence of ACSF or 20 μM YO; n = 3-7 slices from 3-7 mice. (**H**) Traces and summary data of the fluorescence response measured in neurons expressing GRAB_NE1m_ or GRAB_DA1m_. Where indicated, 50 μM NE, 50 μM DA, 20 μM yohimbine (YO), and/or 20 μM haloperidol (Halo) was applied to the cells. n = 3-5 slices from 3-5 mice. The scale bars represent 10 μm. \**p* < 0.05, \*\**p* < 0.01, and \*\*\**p* < 0.001; n.s., not significant (Student’s *t*-test).

NE released into the synapse is recycled back into the presynaptic terminal by the norepinephrine transporter (NET). We therefore tested the sensitivity of GRAB_NE1m_ to NET blockade using desipramine. In the presence of desipramine, electrical stimuli caused a larger fluorescence response in GRAB_NE1m_-expressing neurons compared to ACSF alone (Fig. 4E). Moreover, desipramine significantly slowed the τ_off_ of the fluorescence signal, consistent with reduced reuptake of extracellular NE into the presynaptic terminal. To rule out the possibility that the change in the fluorescence response was caused by a change in synaptic modulation over time, we applied repetitive electrical stimuli at 5-min intervals to GRAB_N_E1m-expressing neurons and found that the stimulation-evoked response was stable for up to 40 min (Fig. 4F). Finally, we examined the specificity of the stimulation-induced response. Compared with a robust response in control conditions, the α-adrenergic antagonist yohimbine blocked the response; moreover, no response was elicited in LC neurons expressing GRAB_N_Emut or in LC neurons expressing a dopamine version of the sensor (GRAB_DA1m_) (Fig. 4G). In contrast, cells expressing GRAB_DA1m_ responded robustly to the application of DA, and the GRAB_NE1m_ and GRAB_DA1m_ responses were abolished by yohimbine and the dopamine receptor antagonist haloperidol, respectively (Fig. 4H). Taken together, these data indicate that GRAB_NE1m_ is both sensitive and specific for detecting endogenous noradrenergic activity in LC neurons.

### GRAB_NE1m_ detects both exogenous NE application and endogenous NE release in awake zebrafish

Zebrafish is both a genetically accessible vertebrate species and an optically transparent organism, thus serving as a suitable model for *in vivo* imaging. We generated the transgenic zebrafish line Tg(HuC:NE1m), which pan-neuronally expresses the GRAB_NE1m_ sensor. Pan-neuronal expression was confirmed by GRAB_NE1m_ fluorescence on the cell membrane of neurons throughout the brain (Fig. 5A). Bath application of 50 μM NE—but not DA at the same concentration—elicited a robust increase in fluorescence intensity that was blocked completely by the subsequent application of 50 μM yohimbine (Fig. 5B-D). In addition, a separate zebrafish line expressing GRAB_NEmut_ did not respond to NE (Fig. 5C,D). Taken together, these data indicate that GRAB_NE1m_ can be used to measure NE in an *in vivo* model.

**Figure 5.**
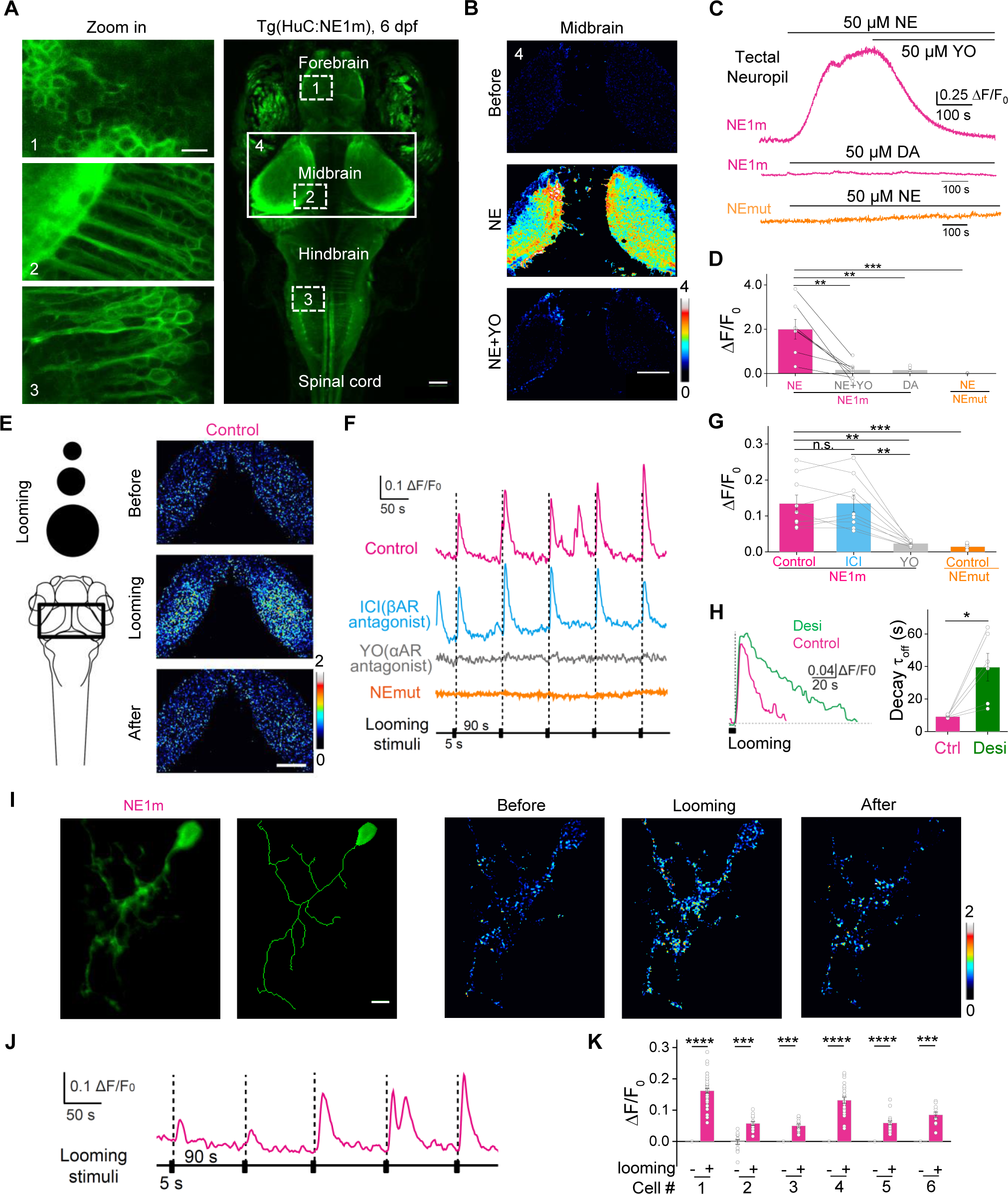
GRAB_NE1m_ can be used to measure noradrenergic activity *in vivo* in transgenic zebrafish. (**A**) *In vivo* confocal image of a Tg(HuC:GRAB_NE1m_) zebrafish expressing GRAB_NE1m_ in neurons driven by the HuC promoter. Larvae at 6 days post-fertilization were used. (**B-D**) Bath application of NE (50 μM) but not DA (50 μM) elicits a significant increase in fluorescence in the tectal neuropil of Tg(HuC:GRAB_NE1m_) zebrafish, but not in GRAB_NEmut_ zebrafish, and this increase is blocked by YO (50 μM), but not ICI 118,551 (50 μM). n = 7. (**E-H**) Visual looming stimuli evoke the release of endogenous NE in the midbrain of GRAB_NE1m_ zebrafish, but not in GRAB_NEmut_ zebrafish. The looming stimuli paradigm is shown in the left of (**E**). Where indicated, YO (50 μM) or ICI 118,551 (50 μM) was applied. Desipramine (Desi, 50 μM) application slowed the decay of looming-induced NE release (**H**). n = 6 for GRAB_NEmut_ and n = 9 for the others. (**I-K**) Single-cell labeling of GRAB_NE1m_ in the midbrain of zebrafish larva (**I**), with looming-evoked responses shown in (**I** and **J**). The summary data for 6 labeled cells are shown in (**K**). The scale bar shown in (**A, left**) represents 10 μm; the scale bars shown in (**A, right**), (**B**) and (**E**) represent 50 μm. The scale bar shown in (**I**) represents 5 μm. \**p* < 0.05, \*\**p* < 0.01, \*\*\**p* < 0.001, and \*\*\*\**p* < 0.0001; n.s., not significant (Wilcoxon matched-pairs signed rank test in panel **H**, all others were analyzed using the paired or unpaired Student’s *t*-test).

Next, we investigated whether GRAB_NE1m_ can be used to measure the dynamics of endogenous noradrenergic activity induced by a visual looming stimulus, which triggers a robust escape response in zebrafish. We applied repetitive looming stimuli while using confocal imaging to measure the fluorescence of GRAB_NE1m_-expressing neurites in the optic tectum (Fig. 5E). Each looming stimulus induced a time-locked increase in GRAB_NE1m_ fluorescence, which was blocked by bath application of yohimbine but was unaffected by the β-adrenergic receptor antagonist ICI 118,551 (Fig. 5F,G). In contrast, the same looming stimuli had no effect in animals expressing GRAB_NEmut_ (Fig. 5F,G). In addition, adding desipramine to block NE reuptake slowed the decay of the fluorescence signal (Fig. 5H). By sparse expression of GRAB_NE1m_ in individual neurons in zebrafish larvae via transient transfection, we were also able to record robust signals corresponding to NE release at single-cell resolution in response to repetitive looming stimuli (Fig. 5I-K), confirming that our GRAB_NE_ sensors can be used to sense NE release at a single-cell level with high spatiotemporal resolution.

### GRAB_NE1m_ detects optogenetically evoked NE release in freely moving mice

Having demonstrated the proof-of-concept in a relatively simple *in vivo* vertebrate system, we next examined whether the GRAB_NE_ sensors can be used to monitor the noradrenergic activity in the mammalian brain by virally expressing GRAB_NE1m_ (non-Cre dependent) together with the optogenetic actuator C1V1 (Cre-dependent) in the LC of Th-Cre mice (Fig. 6A). Optogenetic stimulation of LC NE neurons using 561 nm laser pulses reliably evoked an increase in GRAB_NE1m_ fluorescence in fiber photometry recording of freely moving mice. Moreover, Intraperitoneal (i.p.) injection of desipramine produced a slow progressive increase in basal GRAB_NE1m_ fluorescence (consistent with an increase in extracellular NE levels) and caused an increase in the magnitude and decay time of the light-activated responses. I.p. injection of yohimbine abolished both the increase in basal GRAB_NE1m_ fluorescence and the light-evoked responses (Fig. 6B-D). In contrast, treating mice with either GBR 12909 (a selective blocker of dopamine transporters) or eticlopiride (a specific D2R antagonist) had no effect on the light-evoked responses in GRAB_NE1m_ fluorescence (Fig. 6C-E). To further test the selectivity of GRAB_NE1m_ between NE and dopamine, we co-expressed GRAB_NE1m_ and DIO-C1V1 both in the LC and in the substantia nigra pars compacta (SNc) of Th-Cre mice (Fig. 6F). In these mice, optogenetic stimulation of dopamine neurons in the SNc did not cause any changes in the GRAB_NE1m_ fluorescence in the SNc. In contrast, stimulating NE neurons in the LC produced a clear increase in GRAB_NE1m_ fluorescence (Fig. 6F, G). These results confirm that the increase of GRAB_NE1m_ fluorescence reflects the release of endogenous NE from noradrenergic neurons in the LC.

**Figure 6.**
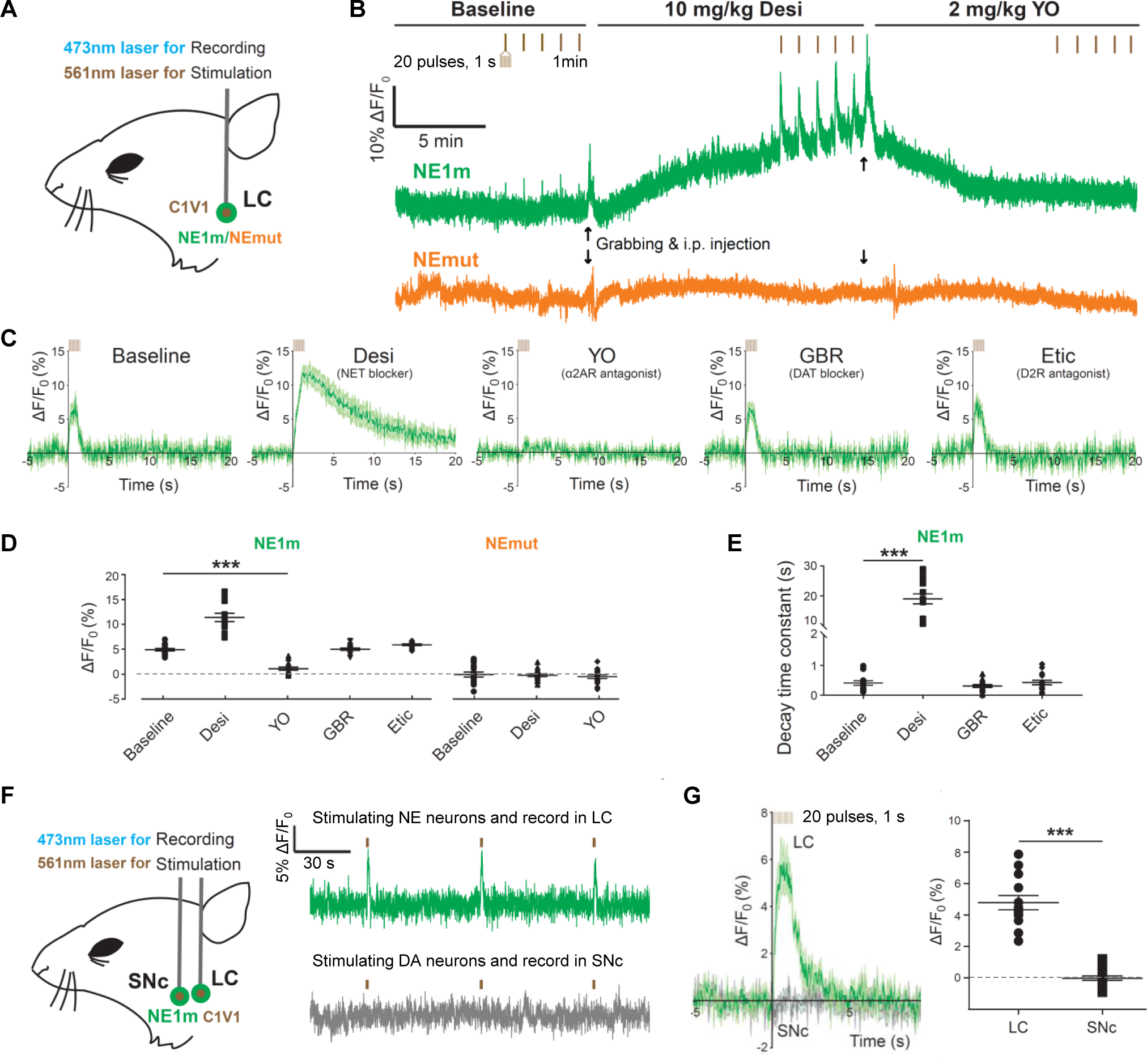
GRAB_NE1m_ can be used to measure optogenetically stimulated noradrenergic activity *in vivo* in freely moving mice. (A) Schematic illustration depicting the experimental design for recording GRAB_NE1m_ and GRAB_NEmut_ fluorescence in response to optical stimulation of C1V1 in the locus coeruleus (LC). (B) Representative traces of optogenetically stimulated GRAB_NE1m_ (top) and GRAB_NEmut_ (bottom) activity in the LC before (baseline, left), 15 min after an i.p. injection of the NET blocker desipramine (10 mg/kg, middle), and 15 min after an i.p. injection of the α2AR antagonist yohimbine (2 mg/kg, right). The vertical tick marks indicate the optogenetic stimuli. Black arrows represent the timing for grabbing and i.p. injection. (**C-D**) Average traces of GRAB_NE1m_ fluorescence (**C**), summary data (**D**), and the decay time constant (**E**) in response to optical stimulation in the LC following treatment with the indicated compounds. n = 15 trials from 3 mice for each condition. (**F,G**) Schematic illustration (**F, left**), representative traces (**F, right**), average fluorescence change (**G, left**), and summary data (**G, right**) for GRAB_NE1m_ in response to optical stimulation of noradrenergic neurons in the LC and dopaminergic neurons in the SNc. \*\*\**p* < 0.001 (for D and E, One-Way ANOVA, for G, Student’s *t*-test).

### Using GRAB_NE1m_ to track endogenous NE dynamics in the mouse hypothalamus during freely moving behaviors

In the brain, the hypothalamus mediates a variety of innate behaviors essential for survival, including feeding, aggression, mating, parenting, and defense (Hashikawa et al., 2016; Sokolowski and Corbin, 2012; Yang and Shah, 2016). The hypothalamus receives extensive noradrenergic projections (Moore and Bloom, 1979; Schwarz and Luo, 2015; Schwarz et al., 2015) and expresses an abundance of α2-adrenergic receptors (Leibowitz, 1970; Leibowitz et al., 1982). Microdialysis studies found that the hypothalamus is among the brain regions that contains high level of NE during stress (McQuade and Stanford, 2000; Pacak et al., 1995; Shekhar et al., 2002; Tanaka, 1999). To better understand the dynamics of NE signaling in the hypothalamus under stress, we virally expressed hSyn-GRAB_NE1m_ in the lateral hypothalamus of C57BL/6 mice. Three weeks after virus injection, we performed fiber photometry recordings of GRAB_NE1m_ fluorescence during a variety of stressful and non-stressful behaviors in freely moving mice (Fig. 7).

**Figure 7.**
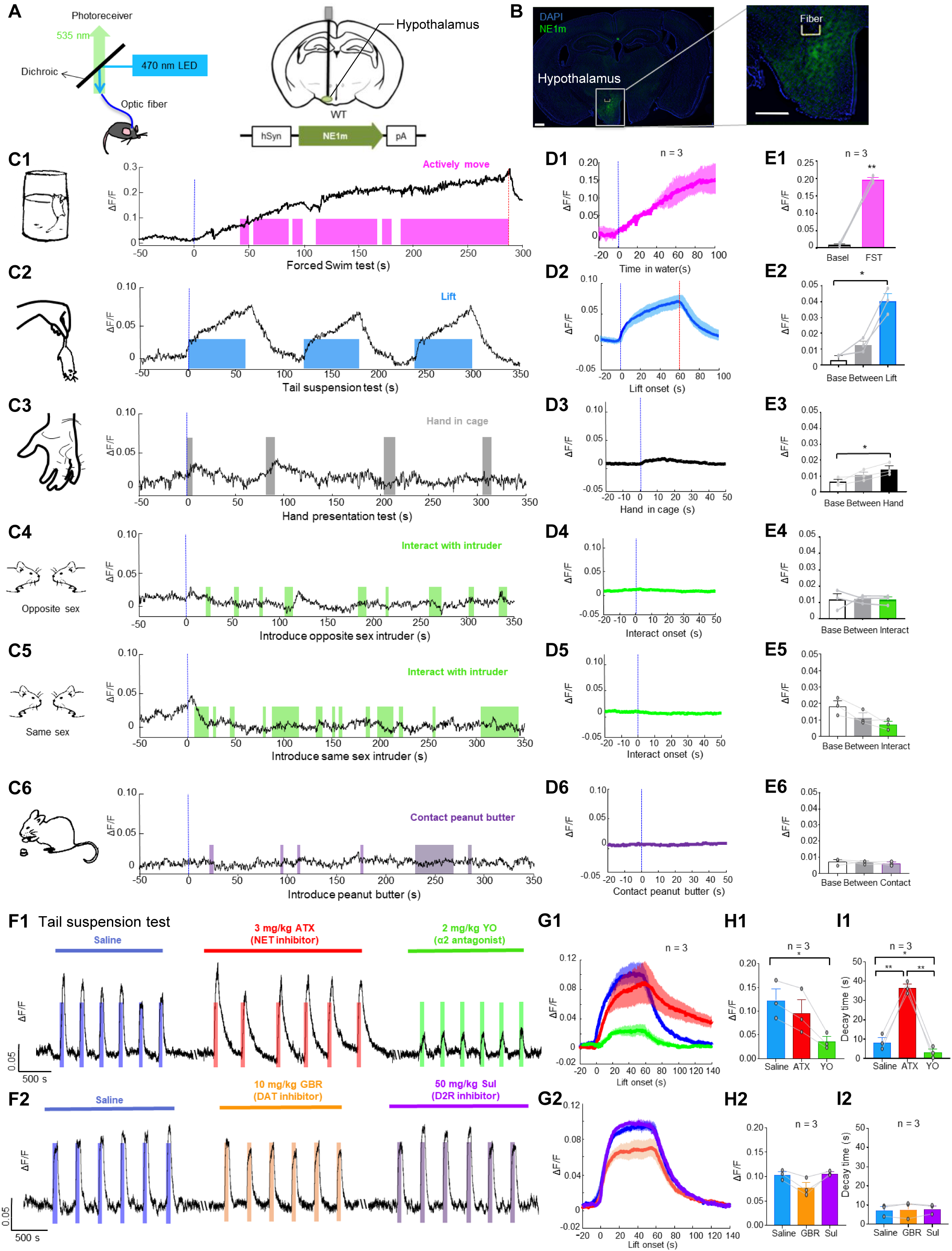
GRAB_NE1m_ can be used to measure noradrenergic activity in the hypothalamus during stress, food-related behavior, and social interaction. (A) Schematic diagrams depicting the fiber photometry recording, virus injection, and recording sites. (B) Histology showing the expression of GRAB_NE1m_ (green) and placement of the recording; the nuclei were counterstained with DAPI (blue). Scale bar: 500μm. (**C1-E6**) Representative traces (**C1-C6**), average per-stimulus histograms (**D1-D6**), and summary data (**E1-E6**) showing normalized GRAB_NE1m_ fluorescence (ΔF/F) before and during the forced swim test (**1**), the tail suspension test (**2**), the hand presentation test (**3**), social interaction with an intruder of the opposite sex (**4**) and the same sex (**5**), and presentation of peanut butter (**6**). n = 3 animals each. (**F**) Representative traces of GRAB_NE1m_ fluorescence during the tail suspension test 10 minutes after saline injection, 25 minutes after atomoxetine (ATX) or yohimbine (YO) injection, and 15 minutes after GBR 12909 or sulpiride (Sul) injection. (**G-I**) Average peri-stimulus histograms (**H**), peak change in GRAB_NE1m_ fluorescence, and post-test decay time measured during the tail suspension test after injection of the indicated compounds. n = 3 each. The Shapiro-Wilk normality test was performed; if the test revealed that the followed a normal distribution, a paired Student’s *t*-test or one-way repeated measures ANOVA followed by Tukey’s multiple comparisons was performed. If the values did not follow a normal distribution, a non-parametric ANOVA (Friedman’s test) was performed followed by Dunn’s multiple comparisons test. In (**C**) and (**D**), the blue dotted lines represent the start of the stimulus, and the red dotted lines represent the end of the trial. \**p* < 0.05 and \*\**p* < 0.01.

During forced swim test and tail suspension test (both of which were stressful), we observed a significant increase in GRAB_NE1m_ fluorescence. During forced swim test, the fluorescence signal increased continuously regardless of the animal’s movements and started to decrease only after the animal was removed from the water (Fig. 7C1-E1). During the 60-s tail suspension test, the signal began to rise when the animal was first pursued by the experimenter’s hand, increased continuously while the animal was suspended by the tail, and decreased rapidly back to baseline levels when the animal was returned to its home cage (Fig. 7C2-E2). Additionally, when a human hand was placed in front of the animal, we observed a small and transient increase in GRAB_NE1m_ fluorescence (Fig. 7C3-E3). In contrast, the presence of a non-aggressive mouse of either the same or the opposite sex or close social interaction with the conspecific (7C4-E4, C5-E5) caused no significant change in GRAB_NE1m_ fluorescence. Lastly, neither sniffing nor eating a food attractant—in this case, peanut butter—had an effect on GRAB_NE1m_ fluorescence (Fig. 7C6-E6). These data provide evidence that noradrenergic activity in the lateral hypothalamus occurs primarily under stressful conditions.

Finally, to confirm that the GRAB_NE1m_ sensor indeed detects changes in NE concentration instead of other monoamine neurotransmitters, such as dopamine, we injected mice with a specific NET inhibitor atomoxetine (3 mg/kg i.p.) to inhibit the reuptake of NE. Although atomoxetine had no effect on the peak change in GRAB_NE1m_ fluorescence during the tail suspension test, it significantly slowed the return to baseline levels after each tail suspension (Fig. 7F1-I1); in contrast, treating mice with the α-adrenergic receptor antagonist yohimbine (2 mg/kg) both decreased the peak change in GRAB_NE1m_ fluorescence and significantly accelerated the return to baseline (Fig. 7F1-I1). Treating mice with either the selective DAT inhibitor GBR 12909 (10 mg/kg, i.p.) or the D2 receptor antagonist sulpiride (50 mg/kg, i.p.) had no effect on the peak change in GRAB_NE1m_ fluorescence or the time to return to baseline (Fig. 7F2-I2). In summary, these data demonstrate that our GRAB_NE_ sensors are suitable for monitoring endogenous noradrenergic activity in real time, with high spatiotemporal precision, during freely moving behavior in mammals.

## Discussion

Here, we report the development and validation of GRAB_NE1m_ and GRAB_NE1h_, two genetically encoded norepinephrine/epinephrine sensors that can be used both *in vitro* and *in vivo* to monitor noradrenergic activity with high temporal and spatial resolution, high ligand specificity, and cell type specificity. In mouse acute brain slices, our GRAB_NE_ sensors detected NE release from the LC in response to electrical stimulation. In zebrafish, the GRAB_NE_ sensors reported looming-induced NE release with single-cell resolution. In mice, the GRAB_NE_ sensors reported the time-locked release of NE in the LC triggered by optogenetic stimulation, as well as changes in hypothalamic NE levels during a variety of stress-related behaviors.

Compared to existing methods for detecting NE, our GRAB_NE_ sensors have several distinct advantages. First, NE has been difficult to distinguish from DA *in vivo* (*e.g.* by fast-scan cyclic voltammetry) (Park et al., 2009; Robinson et al., 2003), largely because of their structural similarities with only one hydroxyl group difference. Our GRAB_NE_ sensors have extremely high *specificity* for NE over other neurotransmitters and chemical modulators, including DA (Figs. 2H, 3F). GRAB_NE1m_ has a roughly 1000-fold higher affinity for NE over DA when expressed in neurons, even better than the 85-fold difference of the wild-type α2-adrenergic receptor. Thus, our GRAB_NE_ sensors provide new opportunities to probe the dynamics of noradrenergic activity with high specificity, which is particularly valuable when studying the many brain regions that receive overlapping dopaminergic and noradrenergic inputs. One thing to note is that GRAB_NE_ sensors are engineered from the α2a receptor, which may not be suitable for pharmacological investigation of α2a receptor related regulations.

Second, our GRAB_NE_ sensors have extremely high *sensitivity* for NE. Specifically, the EC_50_ for NE approaches sub-micromolar levels, with a 200%—or higher—increase in fluorescence intensity upon binding NE. By comparison, recently published FRET-based NE indicators produce a signal change of ≤10% under optimal conditions (Wang et al., 2018a; Wang et al., 2018b). Thus, GRAB_NE_ sensors have much improved characteristics to monitor endogenous *in vivo* NE dynamics. Third, GRAB_NE_ sensors have brightness and photostability properties that rival EGFP, which permits stable recordings across extended experimental sessions. Fourth, because they provide sub-second response kinetics and are genetically encoded, our GRAB_NE_ sensors can non-invasively report noradrenergic activity *in vivo* with single-cell resolution and a high recording rate (∽30 Hz). Finally, because the GRAB_NE_ sensors can traffic to various surface membranes, including the cell body, dendrites, and axons, and because they perform equally well in these membrane compartments, they can provide subcellular spatial resolution, which is essential for understanding compartmental NE signaling *in vivo*.

Ligand binding to endogenous GPCRs drives G-protein activation and receptor internalization. If present in GRAB_NE_ sensors, these responses could interfere with endogenous signaling fidelity and disrupt normal neuronal activity. To assess this risk, we characterized the downstream coupling of our GRAB_NE_ sensors with both G protein–independent and G protein–dependent pathways. Importantly, the introduction of the cpEGFP moiety in the GRAB_NE_ sensors resulted in non-detectable engagement of arrestin-mediated desensitization/internalization, which ensures more consistent surface expression of the sensor and that the GRAB_NE_ sensors do not inadvertently activate arrestin-dependent signaling. With respect to G protein–dependent signaling, we found that although physiological levels of NE robustly induce a change in GRAB_NE1m_ fluorescence, they do not engage downstream G protein signaling (Fig. 2J-M).

Noradrenergic projections throughout the brain originate almost exclusively from the LC, and NE release plays a role in a wide range of behaviors, including cognition and the regulation of arousal, attention, and alertness (Berridge and Waterhouse, 2003; Li et al., 2018; Schwarz et al., 2015). In this respect, it is interesting to note that our *in vivo* experiments revealed that GRAB_NE_ sensors can reliably report looming-evoked NE release in the optic tectum of live zebrafish. Moreover, our fiber photometry recordings of GRAB_NE_ sensors in the hypothalamus of freely behaving mice revealed specific changes in noradrenergic activity under stressful conditions (e.g., a tail lift or forced swimming), whereas non-stressful conditions such as feeding and social interaction did not appear to alter noradrenergic activity. These data are generally consistent with previous data obtained using microdialysis to measure NE (McQuade and Stanford, 2000; Pacak et al., 1995; Shekhar et al., 2002; Tanaka, 1999). Importantly, however, our approach yielded a more temporally precise measurement of noradrenergic activity with the promise of higher spatial and cell-type specificity.

NE circuits of the LC receive heterogeneous inputs from a broad range of brain regions and send heterogeneous outputs to many brain regions (Schwarz et al., 2015). Congruously, altered noradrenergic activity has been associated with a broad range of brain disorders and conditions, including ADHD, PD, depression, and anxiety (Marien et al., 2004). The complexity of these disorders may, in part, reflect the complexities of noradrenergic circuits and signals, which previous tools have been unable to fully dissect. Thus, understanding the regulation and impact of noradrenergic activity during complex behavior demands technological advances, such as the GRAB_NE_ sensors we present here. Deploying these in concert with other cell-specific tools for reporting (Jing et al., 2018; Patriarchi et al., 2018; Sun et al., 2018) and manipulating neurotransmitter levels (Fenno et al., 2011; Urban and Roth, 2015) should increase our understanding of the circuits and mechanisms that underlie brain functions in both health and diseases.

## Experimental model and subject details

### Primary cultures

Rat cortical neurons were prepared from postnatal day 0 (P0) Sprague-Dawley rat pups (both male and female, randomly selected; Beijing Vital River). In brief, cortical neurons were dissociated from dissected P0 rat brains in 0.25% Trypsin-EDTA (Gibco), plated on 12-mm glass coverslips coated with poly-D-lysine (Sigma-Aldrich), and cultured at 37°C in 5% CO_2_ in neurobasal medium (Gibco) containing 2% B-27 supplement, 1% GlutaMax, and 1% penicillin-streptomycin (Gibco).

### Cell lines

HEK293T cells were obtained from ATCC (CRL-3216) and verified based on their morphology under the microscope and by their growth curve. Stable cell lines expressing the wild-type α2-adrenergic receptor or various GRAB_NE_ sensors were constructed by co-transfecting cells with the pPiggyBac plasmid carrying target genes with Tn5 transposase into a stable HEK293T-based cell line expressing chimeric Gαq/i and AP-TGFα (Inoue et al., 2012). Cells that stably expressed the target genes were selected by treating with 2 mg/ml Puromycin (Sigma) after reaching 100% confluence. The HTLA cells used for the TANGO Assay stably express a tTA-dependent luciferase reporter and a β-arrestin2-TEV fusion gene and were a gift from Bryan L. Roth (Kroeze et al., 2015). All cell lines were cultured at 37°C in 5% CO_2_ in DMEM (Gibco) supplemented with 10% (v/v) fetal bovine serum (Gibco) and 1% penicillin-streptomycin (Gibco).

### Mice

All procedures regarding animals were approved by the respective Animal Care and Use Committees at Peking University, New York University, University of Southern California and the US National Institutes of Health, and were performed in compliance with the US National Institutes of Health guidelines for the care and use of laboratory animals. Wild-type Sprague-Dawley rat pups (P0) were used to prepare cultured cortical neurons. Wild-type C57BL/6 and Th-Cre mice (MMRRC_031029-UCD, obtained from MMRRC) were used to prepare the acute brain slices and for the *in vivo* mouse experiments. Experimental Th-Cre mice were produced by breeding Th-Cre hemizygous BAC transgenic mice with C57BL/6J mice. All animals were housed in the animal facility and were family-housed or pair-housed in a temperature-controlled room with a 12hr-12h light-dark cycle (10 pm to 10 am light) with food and water provided *ad libidum*. All *in vivo* mouse experiments were performed using 2-12-month-old mice of both sexes.

### Zebrafish

The background strain for these experiments is the albino strain slc45a2b4. To generate transgenic zebrafish, Both the pTol2-HuC:GRAB_NE1m_ plasmid and Tol2 mRNA were co-injected into single-cell stage zebrafish eggs, and the founders of HuC:NE1m were screened. HuC:NEmut transgenic fish were generated as described above using the pTol2-HuC:GRAB_NEmut_ plasmid. Adult fish and larvae were maintained on a 14h-10h light-dark cycle at 28°C. All experimental larvae were raised to 6-8 days post-fertilization (dpf) in 10% Hank’s solution, which consisted of (in mM): 140 NaCl, 5.4 KCl, 0.25 Na_2_HPO_4_, 0.44 KH_2_PO_4_, 1.3 CaCl_2_, 1.0 MgSO_4_, and 4.2 NaHCO_3_ (pH 7.2). Larval zebrafish do not undergo sex differentiation prior to 1 month post-fertilization (Singleman and Holtzman, 2014).

## Method details

### Molecular cloning

The molecular clones used in this study were generated by Gibson Assembly using DNA fragments amplified using primers (Thermo Fisher Scientific) with 25-bp overlap. The Gibson Assembly cloning enzymes consisted of T5-exonuclease (New England Biolabs), Phusion DNA polymerase (Thermo Fisher Scientific), and Taq ligase (iCloning). Sanger sequencing was performed using the sequencing platform at the School of Life Sciences of Peking University in order to verify the sequence of all clones. All cDNAs encoding the candidate GRAB_NE_ sensors were cloned into the pDisplay vector (Invitrogen) with an upstream IgK leader sequence and a downstream IRES-mCherry-CAAX cassette to label the cell membrane. The cDNAs of select adrenergic receptor candidates were amplified from the human GPCR cDNA library (hORFeome database 8.1), and cpEGFP from GCaMP6s was inserted into the third intracellular loop (ICL3). The insertion sites for the GRAB_NE_ sensors were screened by truncating the ICL3 of the α2-adrenergic receptor at the 10-amino acid (AA) level, followed by fine-tuning at the 1-AA level. Coupling linkers were randomized by PCR amplification using randomized NNB codons in target sites. Other cDNAs used to express the GRAB_NE_ sensors in neurons were cloned into the pAAV vector using the human synapsin promoter (hSyn) or TRE promoter. pAAV-CAG-tTA was used to drive expression of the TRE promoter. The plasmids carrying compartmental markers were cloned by fusing EGFP-CAAX, RFP-CAAX (mScarlet), KDELR-EGFP, PSD95-RFP, and synaptophysin-RFP into the pDest vector. To characterize signaling downstream of the GRAB_NE_ sensors, we cloned the sensors and the wild-type α2-adrenergic receptor into the pTango and pPiggyBac vector, respectively. GRAB_NE1m_-SmBit and α2AR-SmBit constructs were derived from β2AR-SmBit (Wan et al., 2018) using a BamHI site incorporated upstream of the GGSG linker. LgBit-mGsi was a gift from Nevin A. Lambert.

### Expression of GRAB_NE_ sensors in cultured cells and in vivo

The GRAB_NE_ sensors were characterized in HEK293T cells and cultured rat cortical neurons, with the exception of the TANGO assay and TGFα shedding assay. HEK293T cells were passaged with Trypsin-EDTA (0.25%, phenol red; Gibco) and plated on 12-mm size 0 glass coverslips in 24-well plates and grown to ∽70% confluence for transfection. HEK293T cells were transfected by incubating cells with a mixture containing 1 μg of DNA and 3 μg of PEI for 6 h. Imaging was performed 24-48 h after transfection. Cells expressing GRAB_NE_ sensors for screening were plated on 96-well plates (PerkinElmer).

Cultured neurons were transfected using the calcium phosphate method at 7-9 DIV. In brief, the neurons were incubated for 2 h in a mixture containing 125 mM CaCl_2_, HBS (pH 7.04), and 1.5 μg DNAh. The DNA-Ca_3_(PO4)_2_ precipitate was then removed from the cells by washing twice with warm HBS (pH 6.80). Cells were imaged 48 h after transfection.

For *in vivo* expression, the mice were anesthetized by an i.p. injection of 2,2,2-tribromoethanol (Avetin, 500 mg/kg body weight, Sigma-Aldrich), and then placed in a stereotaxic frame for injection of AAVs using a Nanoliter 2000 Injector (WPI) or Nanoject II (Drummond Scientific) microsyringe pump. For the experiments shown in Figures 4 and 6, the AAVs containing hSyn-GRAB_NE1m/NE1mut/DA1m_ and Ef1a-DIO-C1V1-YFP were injected into the LC (AP: −5.45 mm relative to Bregma; ML: ±1.25 mm relative to Bregma; and DV: -2.25 mm from the brain surface) or SNc (AP: -3.1 mm relative to Bregma; ML: ±1.5 mm relative to Bregma; and DV: -3.8 mm from the brain surface) of wild-type or Th-Cre mice. For the experiments shown in Figure 7, 100 nl of AAV9-hSyn-GRAB_NE1m_ (Vigene, 1×10^13^ titer genomic copies per ml) were unilaterally into the hypothalamus (AP: -1.7 mm relative to Bregma; ML: 0.90 mm relative to Bregma; and DV: -6.05 mm from the brain surface) of wild-type (C57BL/6) mice at a rate of 10 nl/min.

### Fluorescence imaging of HEK293T cells and cultured neurons

HEK293T cells and cultured neurons expressing GRAB_NE_ sensors were screened using an Opera Phenix high-content imaging system (PerkinElmer) and imaged using an inverted Ti-E A1 confocal microscope (Nikon). A 60x/1.15 NA water-immersion objective was mounted on the Opera Phenix and used to screen GRAB_NE_ sensors with a 488-nm laser and a 561-nm laser. A 525/50 nm and a 600/30 nm emission filter were used to collect the GFP and RFP signals, respectively. HEK293T cells expressing GRAB_NE_ sensors were first bathed in Tyrode’s solution and imaged before and after addition of the indicated drugs at the indicated concentrations. The change in fluorescence intensity of the GRAB_NE_ sensors was calculated using the change in the GFP/RFP ratio. For confocal microscopy, the microscope was equipped with a 40x/1.35 NA oil-immersion objective, a 488-nm laser, and a 561-nm laser. A 525/50 nm and a 595/50 nm emission filter were used to collect the GFP and RFP signals, respectively. GRAB_NE_-expressing HEK293T cells and neurons were perfused with Tyrode’s solutions containing the drug of interest in the imaging chamber. The photostability of GRAB_NE_ sensors and EGFP was measured using a confocal microscope (for 1-photon illumination) equipped with a 488-nm laser at a power setting of ∽350 μW, and using a FV1000MPE 2-photon microscope (Olympus, 2-photon illumination) equipped with a 920-nm laser at a power setting of ∽27.5 mW. The illuminated region was the entire HEK293T cell expressing the target protein, with an area of ∽200 μm^2^. Photolysis of NPEC-caged-NE (Tocris) was performed by combining fast scanning with a 76-ms pulse of 405-nm laser illumination by a confocal microscope.

### TANGO assay

NE at various concentrations (ranging from 0.1 nM to 100 μM) was applied to α2AR-expressing or NE1m-/NE1h-expressing HTLA cells (Kroeze et al., 2015). The cells were then cultured for 12 hours to allow expression of the luciferase gene. Furimazine (NanoLuc Luciferase Assay, Promega) was then applied to a final concentration of 5 μM, and luminescence was measured using a VICTOR X5 multilabel plate reader (PerkinElmer).

### TGFα shedding assay

Stable cell lines expressing Gαi-AP-TGFα together with the wild-type α2AR or GRAB_NE_ sensors were plated in a 96-well plate and treated by the addition of 10 μl of a 10x solution of NE in each well, yielding a final NE concentration ranging from 0.1 nM to 100 μM. Absorbance at 405 nm was read using a VICTOR X5 multilabel plate reader (PerkinElmer). TGFα release was calculated as described previously (Inoue et al., 2012). Relative levels of G protein activation were calculated as the TGFα release of GRAB_NE_ sensors normalized to the release mediated by wild-type α2AR.

### FSCV

Fast-scan cyclic voltammetry was performed using 7-μm carbon fiber microelectrodes. Voltammograms were measured with a triangular potential waveform from -0.4 V to +1.1 V at a scan rate of 400 V/s and a 100-ms interval. The carbon fiber microelectrode was held at -0.4 V between scans. Voltammograms measured in the presence of various different drugs in Tyrode’s solution were generated using the average of 200 scans followed by the subtraction of the average of 200 background scans. Currents were recorded using the Pinnacle tethered FSCV system (Pinnacle Technology). Pseudocolor plots were generated using Pinnacle FSCV software. The data were processed using Excel (Microsoft) and plotted using Origin Pro (OriginLab).

### Luciferase complementation assay

The luciferase complementation assay was performed as previously described (Wan et al., 2018). In brief, ∽48h after transfection the cells were washed with PBS, harvested by trituration, and transferred to opaque 96-well plates containing diluted NE solutions. Furimazine (Nano-Glo; 1:1000; Promega) was added to each well immediately prior to performing the measurements with Nluc.

### Fluorescence imaging of GRAB_NE_ in brain slices

Fluorescence imaging of acute brain slices was performed as previously described (Sun et al., 2018). In brief, the animals were anesthetized with Avertin, and acute brain slices containing the LC region or the hippocampus region were prepared in cold slicing buffer containing (in mM): 110 choline-Cl, 2.5 KCl, 1.25 NaH_2_PO_4_, 25 NaHCO_3_, 7 MgCl_2_, 25 glucose, and 2 CaCl_2_. Slices were allowed to recover at 35°C in oxygenated Ringers solution containing (in mM): 125 NaCl, 2.5 KCl, 1.25 NaH_2_PO_4_, 25 NaHCO_3_, 1.3 MgCl_2_, 25 glucose, and 2 CaCl_2_ for at least 40 minutes before experiments. An Olympus FV1000MPE two-photon microscope equipped with a 40x/0.80 NA water-immersion objective and a mode-locked Mai Tai Ti:Sapphire laser (Spectra-Physics) tuned to 920 nm were used for imaging the slices. For electrical stimulation, a concentric electrode (model #CBAEC75, FHC) was positioned near the LC region, and the imaging and stimuli were synchronized using an Arduino board controlled using a custom-written program. The imaging speed was set at 0.148 s/frame with 128 x 96 pixels in each frame. The stimulation voltage was set at ∽6 V, and the duration of each stimulation was typically 1 ms. Drugs were either delivered via the perfusion system or directly bath-applied in the imaging chamber.

For immunostaining of brain sections, GRAB_NE_-expressing mice were anesthetized with Avetin, and the heart was perfused with 0.9% NaCl followed by 4% paraformaldehyde (PFA). The brain was then removed, placed in 4% PFA for 4 h, and then cryoprotected in 30% (w/v) sucrose for 24 h. The brain was embedded in tissue-freezing medium, and 50-µm thick coronal sections were cut using a Leica CM1900 cryostat (Leica, Germany). A chicken anti-GFP antibody (1:500, Abcam, #ab13970) was used to label GRAB_NE_, and a rabbit anti-DBH antibody (1:50, Abcam, #ab209487) was used to label adrenergic terminals in the hippocampus. Alexa-488-conjugated goat-anti-chicken and Alexa-555-conjugated goat-anti-rabbit secondary antibodies were used as the secondary antibody, and the nuclei were counterstained with DAPI. The sections were imaged using a confocal microscope (Nikon).

### Fluorescence imaging of zebrafish

Tg(HuC:GRAB-NE_1m_) zebrafish larvae were imaged by using an upright confocal microscope (Olympus FV1000, Japan) equipped with a 20x water-dipping objective (0.95 NA). The larvae were first paralyzed with α-bungarotoxin (100 μg/ml, Sigma), mounted dorsal side up in 1.5% low melting-point agarose (Sigma), and then perfused with an extracellular solution consisting of (in mM) 134 NaCl, 2.9 KCl, 4 CaCl_2_, 10 HEPES, and 10 glucose (290 mOsmol/L, pH 7.8). Images were acquired at 1-2 Hz with a view field of 800 × 800 pixels and a voxel size was 0.62 × 0.62 × 2.0 μm^3^ (x × y × z). To detect the sensor’s response to exogenous NE, 50 μM L-(-)-norepinephrine (+)-bitartrate salt monohydrate (Sigma) in 5 μM L-ascorbic acid and 50 μM yohimbine hydrochloride (TOCRIS) were sequentially applied to the bath. To detect endogenous NE release, visual looming stimuli, which mimic approaching objects or predators (Yao et al., 2016) were projected to the larvae under a red background. Each trial lasted 5 s, and 5 trials were performed in a block, with a 90-s interval between trials. To examine the specificity of responses, ICI 118,551 hydrochloride (50 μM, Sigma) and desipramine hydrochloride (50 μM, Sigma) were applied. Looming stimuli in transiently transfected HuC:GRAB_NE1m_ zebrafish were measured at single-cell resolution by using the same conditions described above.

### Fiber photometry recording in freely moving mice during optical stimulation

In the all-optic experiments shown in Figure 6, multimode optical fiber probes (105/125 µm core/cladding) were implanted into the LC (AP: -5.45 mm relative to Bregma; ML: ±0.85 mm relative to Bregma; and DV: -3.5 mm from the brain surface) and the SNc (AP: -3.1 mm relative to Bregma; ML: ±1.5 mm relative to Bregma; and DV: -3.85 mm from the brain surface) in mice four weeks after viral injection. Fiber photometry recording in the LC and/or SNc was performed using a 473-nm laser with an output power of 25 µW measured at the end of the fiber. The measured emission spectra were fitted using a linear unmixing algorithm (https://www.niehs.nih.gov/research/atniehs/labs/ln/pi/iv/tools/index.cfm). The coefficients generated by the unmixing algorithm were used to represent the fluorescence intensities of various fluorophores (Meng et al., 2018). To evoke C1V1-mediated NE/DA release, pulse trains (10-ms pulses at 20 Hz for 1 s) were delivered to the LC/SNc using a 561-nm laser with an output power of 9.9 mW measured at the end of the fiber.

### Fiber photometry recording in mice during behavioral testing

For the experiments in Figure 7, a fiber photometry recording set-up was generated and used as previously described (Falkner et al., 2016). GRAB_NE1m_ was injected into the lateral hypothalamus (Bregma AP: -1.7mm; ML: 0.90 mm DV: -4.80 mm) of C57BL/6 mice in a volume of 100 nl containing AAV9-hSyn-GRAB_NE1m_ (Vigene, 1×10^13^ titer genomic copies per ml) at 10 nl/min. A 400-µm optic fiber (Thorlabs, BFH48-400) housed in a ceramic ferrule (Thorlabs, SFLC440-10) was implanted 0.2 mm above the injection site. The virus was left to incubate for three weeks. Prior to fiber photometry recording, a ferrule sleeve was used to connect a matching optic fiber to the implanted fiber. For recordings, a 400Hz sinusoidal blue LED light (30 µW; M470F1 driven by an LEDD1B driver; both from Thorlabs) was bandpass-filtered (passing band: 472 ± 15 nm, Semrock, FF02-472/30-25) and delivered to the brain in order to excite GRAB_NE1m_. The emission light passed through the same optic fiber, through a bandpass filter (passing band: 534 ± 25 nm, Semrock, FF01-535/50), and into a Femtowatt Silicon Photoreceiver, which recorded the GRAB_NE1m_ emission using an RZ5 real-time processor (Tucker-Davis Technologies). The 400-Hz signals from the photoreceiver were extracted in real time using a custom program (Tucker-Davis Technologies) and used to reflect the intensity of the GRAB_NE1m_ fluorescence signal.

### Behavioral assays

All behavioral tests were performed at least one hour after the onset of the dark cycle. For the tail suspension test, each mouse was gripped by the tail and lifted off the bottom of its cage six times for 60 s each, with at least one minute between each lift. For the forced swim test, the mouse was gently placed in a 1000-ml conical flask containing lukewarm water and removed after 4-6 minutes. After removal from the water, the mouse was gently dried with paper towels and placed in the home cage on a heating pad. For conspecific assays, an adult C57BL/6 group-housed mouse of either sex was placed inside the test mouse’s cage for 10 minutes. No sexual behavior or aggressive behavior was observed during the interaction. For the food assay, ∽4g of peanut butter was placed in the cap of a 15-ml plastic tube and placed inside of the test mouse’s cage for 10 minutes. During that period, the test mouse was free to explore, sniff, and eat the peanut butter. All videos were acquired at 25 frames per second and manually annotated frame-by-frame using a custom MATLAB program (Lin et al., 2011). “Contact” with the social stimulus refers to the period in which the test mouse sniffed or was sniffed by the intruder. “Contact” with the peanut butter refers to the period in which the test mouse sniffed or ate the peanut butter. “Lift” refers to the period in which the experimenter gripped the mouse’s tail and lifted the mouse into the air.

### Quantification and statistical analysis

For the imaging experiments using cultured HEK293T cells, primary neurons, and brain slices, images were first imported to ImageJ software (National Institutes of Health) for fluorescence intensity readouts, and then analyzed using MATLAB (MathWorks) with a custom-written script or Origin Pro (OriginLab). The fluorescence response traces in the brain slices shown in Figure 4 were processed with 3x binning and then plotted.

Time-lapse images of the zebrafish were analyzed using Fiji to acquire the fluorescence intensity in the region of interest (ROI) in each frame. A custom-written MATLAB program was then used to calculate the change in fluorescence intensity (ΔF/F_0_) as follows: ΔF/F_0_=(F_t_-F_0_)/F_0_, where F_t_ was the fluorescence intensity at time t and F_0_ was the average fluorescence intensity during the entire time window. Statistical analyses were performed using GraphPad Prism 6 and Origin Pro (OriginLab).

For the fiber photometry data shown in Figure 7, the MATLAB function “msbackadj” with a moving window of 25% of the total recording duration was first applied to obtain the instantaneous baseline signal (F_baseline_). The instantaneous ΔF/F was calculated as (F_raw_ – F_baseline_)/F_baseline_, and a peri-stimulus histogram (PSTH) was calculated by aligning the ΔF/F signal of each trial to the onset of the behavior of interest. The response elicited during a behavior was calculated as the average ΔF/F during all trials of a given behavior. The response between behavioral periods was calculated as the average ΔF/F between two behavioral episodes excluding 4 s immediately before the behavior’s onset, as some uncontrolled and/or unintended events (e.g., chasing the animal before the tail suspension test) may have occurred during that period. The baseline signal was calculated as the average ΔF/F 100 s prior to the start of the behavioral test. The peak response after each drug injection was calculated as the average maximum ΔF/F during all tail suspension trials. The decay time was calculated as the average time required to reach half of the peak response.

Except where indicated otherwise, group differences were analyzed using the Student’s *t*-test, Wilcoxon matched-pairs signed rank test, Shapiro-Wilk normality test, one-way ANOVA test, or Friedman’s test. Except where indicated otherwise, all summary data are presented as the mean ± SEM.

## Data and software availability

The custom MATLAB programs using in this study will be provided upon request to the corresponding author.

---

**Figure S1.**
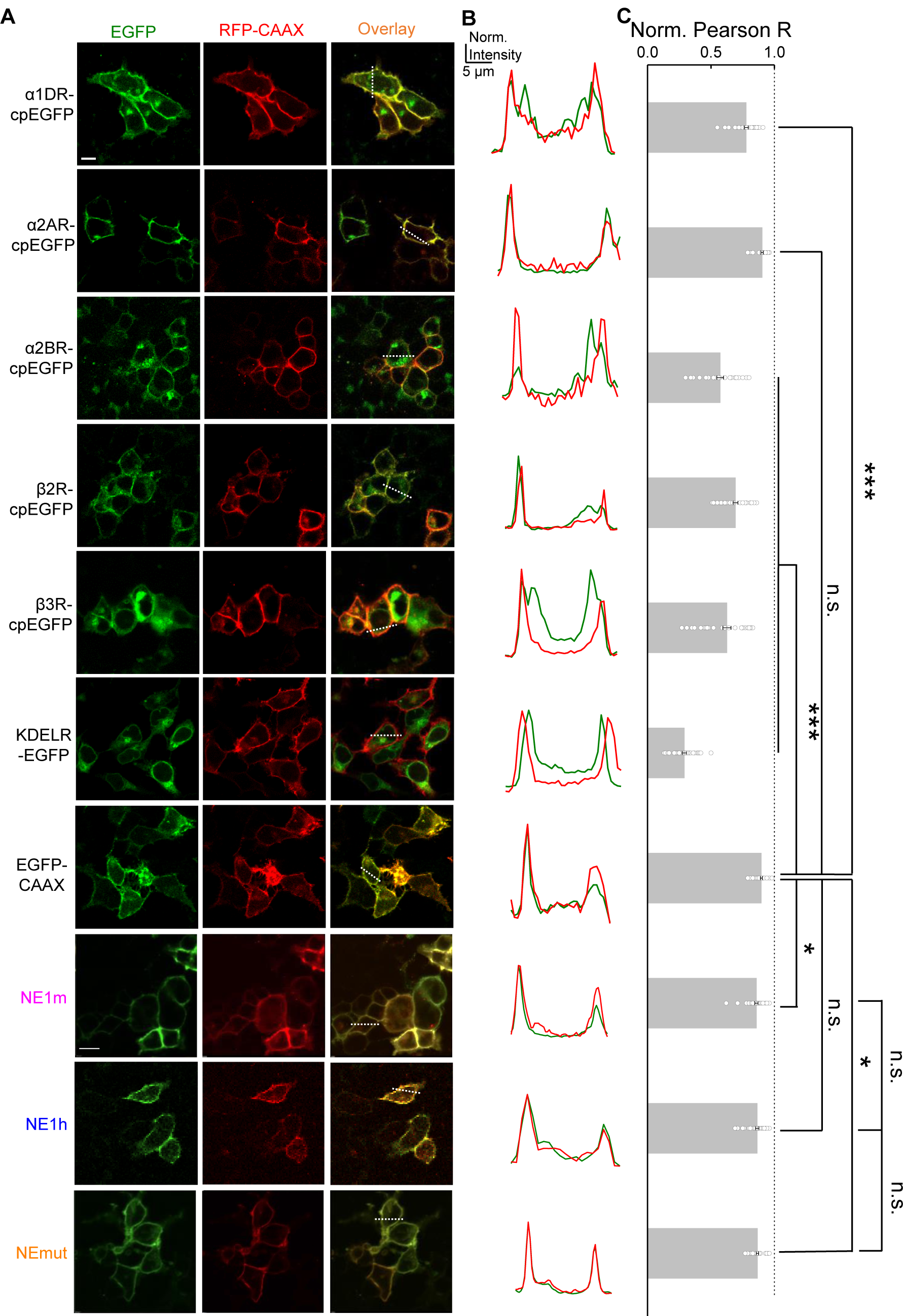
Characterization of the membrane trafficking of a panel of screening candidates (related to Fig. 1). Representative images (**A**) of HEK293T cells co-transfected with the indicated screening candidates (green) together with RFP-CAAX (red) to label the plasma membrane. KDELR-EGFP was used as an ER marker. The dashed white lines indicate the line used for the line-scanning data shown in (**B**) and summarized in (**C**) n = 30 cells from 4-5 cultures. The scale bars in (**A**) represent 10 μm. \**p* < 0.05 and \*\*\**p* < 0.001; n.s., not significant (Student’s *t*-test).

**Figure S2.**
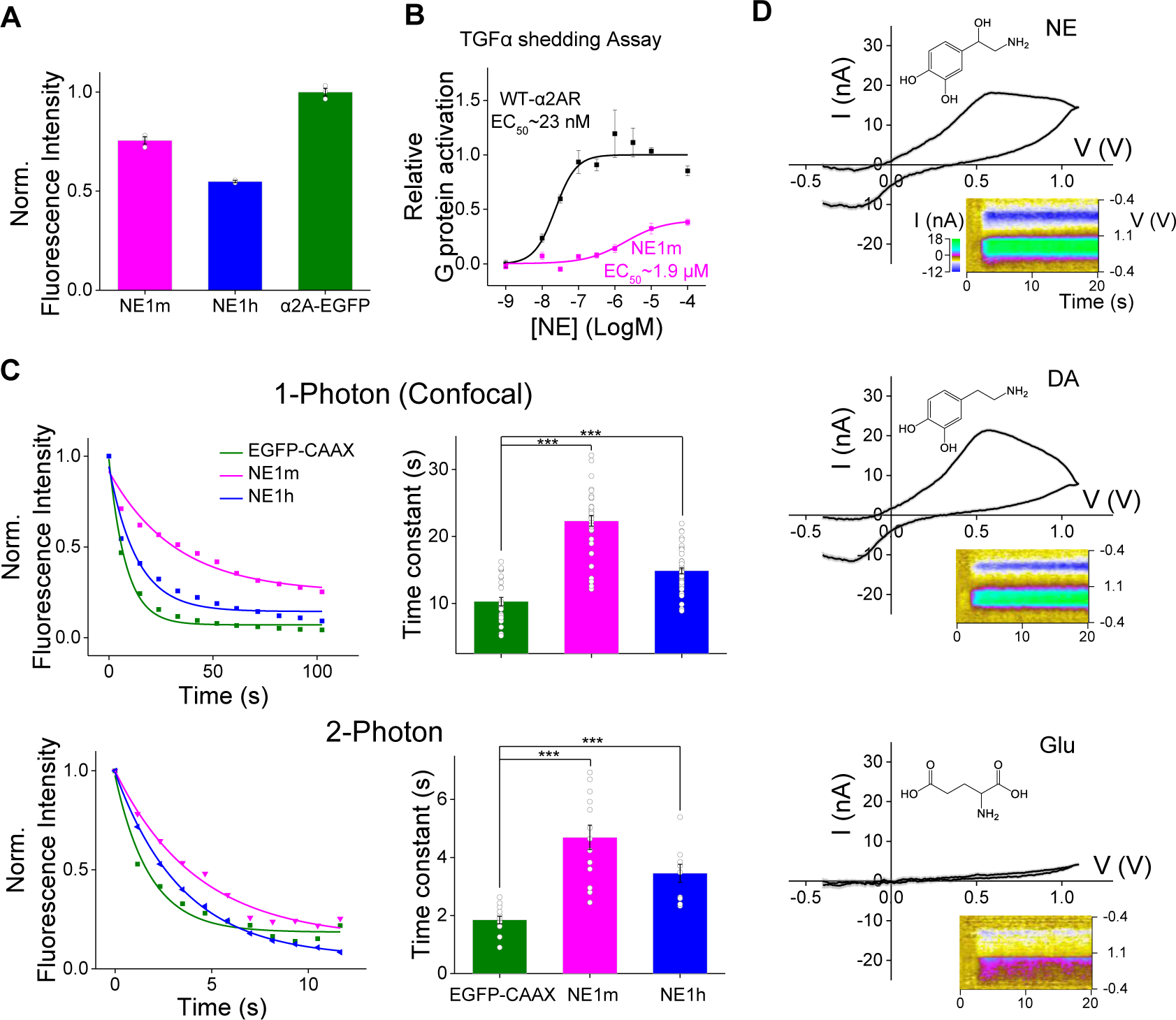
Further characterization of GRAB_NE_ sensors (related to Fig. 2). (A) Fluorescence intensity of GRAB_NE1m_ and GRAB_NE1h_ expressed relative to EGFP-α2AR. n ≥ 2 wells with 300-500 cells per well. (B) G protein activation mediated by GRAB_NE1m_ and wild-type α2AR was measured using the TGFα shedding assay and is expressed relative to α2AR. n = 4 wells with ≥10^5^ cells per well. (**C**) Exemplar (**left**) and summary data (**right**) showing the photostability of GRAB_NE_ sensors and EGFP-CAAX using confocal (**top**) and 2-photon (**bottom**) microscopy. n > 10 cells from at least 3 cultures. (**D**) Exemplar cyclic voltammograms for 10 μM NE (**top**), 10 μM DA (**middle**), and 10 μM Glu (**bottom**) measured using FSCV are shown. The traces were averaged from separate 200 trials. \*\*\**p* < 0.001 (Student’s *t*-test).

**Figure S3.**
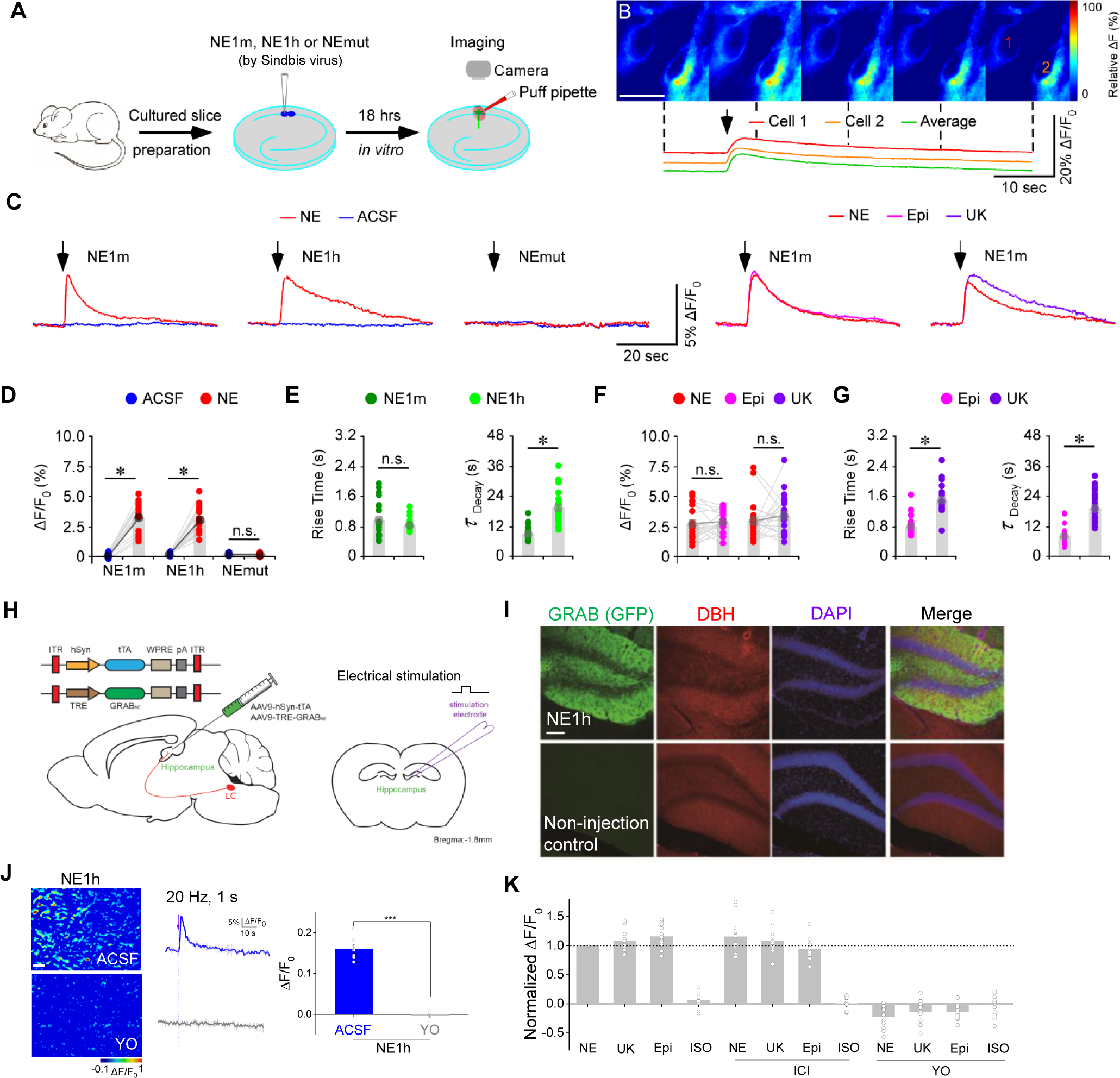
GRAB_NE_ sensors respond selectively to noradrenergic agonists in brain slices (related to Fig. 4). (**A**) Schematic drawing showing the experimental design for measuring CA1 pyramidal neurons in cultured rat hippocampal slices. (**B**) Heat-map images of the change in fluorescence in GRAB_NE1m_-expressing CA1 neurons in response to a 10-ms local application of NE (20 µM). The red and orange traces show the fluorescence responses of two neurons, and the green trace shows the average response of all neurons in the field. The scale represents 20 μm. (C) Fluorescence responses measured in GRAB_NE1m_-, GRAB_NE1h_-, and GRAB_NEmut_-expressing CA1 neurons following a 10-ms puff (arrow) of ACSF, NE (20 µM), Epi (100 µM), or brimonidine (UK, 20 µM). (D) Maximum ΔF/F_0_ response measured in GRAB_NE1m_-, GRAB_NE1h_-, and GRAB_NEmut_-expressing CA1 neurons following a 10-ms puff of ACSF or NE. n = 20-21 cells from 8 animals per group. (**E**) Rise times and decay time constants were measured in CA1 neurons expressing GRAB_NE1m_- and GRAB_NE1h_-expressing CA1 neurons in response to a puff of NE. n = 21 cells from 8 animals. (**F**) Maximum ΔF/F_0_ response measured in GRAB_NE1m_-expressing CA1 neurons following a puff of NE, Epi, or brimonidine (UK). n = 20-21 cells from 8 animals per group. (**G**) Rise times and decay time constants were measured in GRAB_NE1m_-expressing CA1 neurons following a puffs of Epi or brimonidine (UK). (**H**) Schematic illustration depicting AAV-mediated delivery of GRAB_NE1h_ in the mouse hippocampus and bath application of various agonists in the dentate gyrus. (**I**) Example images showing GRAB_NE1h_ (green) expression and dopamine beta hydroxylase (DBH) immunostaining (red) in the dentate gyrus of AAV-GRAB_NE1h_- and control-injected hippocampi. The nuclei were counterstained with DAPI. The scale bar represents 100 μm. (**J**) Electrical stimulation evokes NE release in the hippocampus measured as a change in GRAB_NE1h_ fluorescence. The response was blocked by batch application of yohimbine (YO). Exemplar images (**left**), representative traces (**middle**), and the summary data (**right**) are shown. (**K**) Normalized change in GRAB_NE1h_ fluorescence in response to bath application of the indicated noradrenergic agonists in the presence or absence of ICI 118,551 or yohimbine. The scale bar shown in (**B**) represents 20 μm; the scale bar shown in (**I**) represents 100 μm. The scale bar shown in (**J**) represents 10 μm. \**p* < 0.05 and \*\*\**p* < 0.001; n.s., not significant (Student’s *t*-test, Wilcoxon test, or Mann-Whitney rank sum test).

Author Contributions
Y. L conceived and supervised the project. J.F., M.J., H.Wang, A.D., and Z.W. performed experiments related to sensor development, optimization, and characterization in culture HEK cells, culture neurons and brain slices. Y.Z., P.Z. and J.J.Z designed and performed experiments using Sindbis virus in slices. C.Z., W.C., and J.D. designed and performed experiments on transgenic fish. J.L., J.Zhou, H.Wu, J.,Zou, S.A.H., G.C., and D.L. designed and performed experiments in behaving mice. All authors contributed to data interpretation and data analysis. Y. L and J.F. wrote the manuscript with input from M.J., J.L., and D.L. and help from other authors.

## Acknowledgements

This work was supported by the National Basic Research Program of China (973 Program; grant 2015CB856402), the General Program of National Natural Science Foundation of China (project 31671118), the NIH BRAIN Initiative grant U01NS103558, the Junior Thousand Talents Program of China, the grants from the Peking-Tsinghua Center for Life Sciences, and the State Key Laboratory of Membrane Biology at Peking University School of Life Sciences to Y. L; the Key Research Program of Frontier Sciences (QYZDY-SSW-SMC028) of Chinese Academy of Sciences, and Shanghai Science and Technology Committee (18JC1410100) to J.D.; the NIH grants R01MH101377 and R21HD090563 and an Irma T. Hirschl Career Scientist Award to D.L.; and the Intramural Research Program of the NIH/NIEHS of the United States (1ZIAES103310) to G.C.

We thank Yi Rao for sharing the two-photon microscope and Xiaoguang Lei for the platform support of the Opera Phenix high-content screening system at PKU-CLS. We thank the Core Facilities at the School of Life Sciences, Peking University for technical assistance. We thank Bryan L. Roth and Nevin A. Lambert for sharing stable cell lines and plasmids. We thank Yue Sun, Sunlei Pan, Lun Yang, Haohong Li for inputs on sensors’ characterization and application. We thank Yanhua Huang, Liqun Luo and Mickey London for valuable feedback of the manuscript.

